# Pharmacological co-targeting of Hippo and Wnt pathways induces robust Müller glia proliferation and reprogramming in the adult mouse retina

**DOI:** 10.64898/2026.11.18.767564

**Authors:** Wanling He, Yaqiong Yang, Ziyi Sha, Qinrui Hu, Xiying Zhai, Jintao Shi, Lifan Yang, Zihan Guo, Mouxin Zhang, Yuehua Zhang, Wei Li, Liying Tang, Yi Liao

## Abstract

Reprogramming of endogenous Müller glia (MG) into functional neurons represents a promising therapeutic strategy for the treatment of irreversible retinal blinding diseases. In adult mammals, however, insufficient MG proliferation constitutes a primary bottleneck that restricts successful MG reprogramming and impairs retinal structural and functional restoration. Here, we demonstrated that simultaneous chemical inhibition of Hippo signaling with TRULI and activation of Wnt signaling with CHIR-99021 elicited robust proliferation of endogenous MG in the adult mouse retinal explants. Following combinatorial drug treatment, MG exhibited gradual induction of the retinal progenitor marker PAX6 and the bipolar/photoreceptor lineage determinant Otx2, without concomitant elevation of the fibrotic marker GFAP, supporting the initiation of MG reprogramming. Mechanistically, our findings revealed that YAP and β-catenin might act cooperatively to induce the expression of MYC, a core transcription factor governing cell cycle progression and somatic cell reprogramming. Elevated MYC accumulation in MG transcriptionally activated Cyclin D1 expression, which, together with the downregulation of the cell cycle inhibitor p27^Kip1^, drove efficient MG cell cycle re-entry and proliferation. Collectively, this study establishes a pure chemical combinatorial strategy to trigger robust proliferation and initiate reprogramming of adult mammalian MG. Our findings provide a feasible alternative approach to promote retinal regeneration and lay a foundation for the future clinical treatment of degenerative retinal diseases.

## INTRODUCTION

Retinal degenerative diseases are characterized by structural and functional deterioration of the retina, stemming from acute or progressive loss of key retinal neurons including photoreceptors and retinal ganglion cells. This pathological process ultimately causes irreversible vision loss, and no effective therapeutic interventions currently available for these blinding disorders. Similar to other neurodegenerative diseases of the central nervous system, neuronal loss represents the core pathological hallmark of retinal degeneration. Accordingly, strategies to replenish lost retinal neurons remain a major bottleneck for achieving functional retinal repair.

In mammals, retinal neurons are terminally differentiated and thus incapable of regeneration in adulthood. In contrast, cold-blooded vertebrates such as zebrafish possess a robust retinal regenerative capacity, wherein retinal Müller glia (MG) can be activated upon retinal injury. Activated MG sequentially undergo reprogramming into a progenitor-like state, proliferate and ultimately differentiate to regenerate functional neurons[1]. Although mammalian MG have lost the intrinsic capacity for neuronal regeneration, earlier studies have demonstrated that acute retinal injury can trigger transient upregulation of *cyclin* genes[2] and the progenitor marker gene *Pax6*[3], suggesting that mammalian MG possess latent regenerative potential. Multiple studies have attempted to unlock this regenerative capacity by mimicking zebrafish regenerative mechanisms. For instance, forced expression of the proneural transcription factor Ascl1 combined with epigenetic modulation via the histone deacetylase inhibitor trichostatin A (TSA) induces the transdifferentiation of adult mouse MG into neuron-like cells expressing retinal neuron markers Otx2 and HuC/D[4]. Subsequently, the same group further demonstrated that combinatorial overexpression of developmental transcription factors (Islet1, Pou4f2, Atoh1, and Ascl1) with TSA treatment effectively promotes MG trans-differentiation into retinal ganglion cell-like neurons in injured adult mouse retinas[5]. Parallel studies have shown that β-catenin overexpression stimulates MG proliferation, and subsequent delivery of Otx2, Crx, and Nrl enables MG to differentiate into rod photoreceptors [6]. More recently, genetic suppression of Notch signaling via *Notch1/2* or *Rbpj* ablation has been shown to reprogram endogenous MG into bipolar and amacrine-like neurons in adult mice[7, 8]. Collectively, these findings strongly demonstrate that mammalian MG harbor considerable potentials for retinal neuronal regeneration.

As the principal glial cell type in the mammalian retina, MG are essential for maintaining retinal homeostasis and tissue integrity. MG span the entire retinal thickness and ensheath neuronal somata and processes to sustain structural stability, while also providing critical metabolic support for retinal neurons and their function[9]. Given these indispensable roles, MG loss leads to retinal dysplasia and subsequent degenerative progression[2, 10]. In this regard, direct reprogramming of resident MG into neurons without prior proliferation may compromise retinal structural integrity and tissue homeostasis. Therefore, inducing robust MG proliferation to generate a sufficient cellular reservoir is a prerequisite for safe and effective retinal regeneration. Early mechanistic investigations confirmed that retinal injury enables MG to re-enter the cell cycle via transient Cyclin D3 upregulation and p27^Kip1^ downregulation[2]. However, this endogenous cell cycle response is insufficient to sustain substantial MG proliferation, as MG rapidly exit the cell cycle and undergo reactive gliosis characterized by elevated GFAP expression[2]. To enhance MG proliferative capacity, multiple regulatory strategies have been explored. Intravitreal delivery of growth factors, including EGF, FGF, or FGF combined with insulin, can effectively trigger MG proliferation in injured mouse retinas[11]. Mechanistically, Hippo signaling imposes a critical barrier to cell cycle re-entry in adult murine MG by phosphorylating YAP and thereby suppressing YAP activity. Accordingly, transgenic overexpression of constitutively active YAP5SA to bypass this Hippo-dependent repression promotes spontaneous MG proliferation through upregulation of Cyclin D1[12, 13]. On the other hand, adeno-associated virus (AAV)-mediated β-catenin (downstream effector of Wnt pathway) overexpression potently promotes MG mitotic division in adult mice [6]. Furthermore, AAV-mediated simultaneous Cyclin D1 overexpression and p27^Kip1^ knockdown is sufficient to induce MG proliferation even in intact mouse retinas[14]. Taken together, adult mammalian MG retain intrinsic proliferative competence, which can be efficiently activated via diverse genetic and pharmacological interventions.

Despite the remarkable efficacy of genetic strategies in facilitating MG reprogramming, such approaches raise safety concerns for translational applications[15]. As a promising alternative, small-molecule-based chemical modulation has emerged as a feasible strategy to manipulate cell fate for regenerative purposes[16]. In this context, the present study aimed to investigate whether chemical intervention can efficiently drive MG reprogramming to promote retinal regeneration. Our previous work identified DMX-5804, a selective small-molecule MAP4Ks inhibitor, as a suppressor of Hippo signaling that activates YAP and induces MG proliferation and reprogramming in adult mice[17]. Nevertheless, this single-molecule treatment yields fewer proliferative MG compared with established genetic manipulations[6, 12–14], which severely restricts the efficiency of neuronal regeneration and functional retinal repair. To address this limitation, the current study demonstrates that combined treatment with the Hippo inhibitor TRULI and the Wnt activator CHIR-99021 robustly enhanced MG proliferation. These proliferating MG transiently upregulated the retinal progenitor marker PAX6 and subsequently expressed the bipolar and photoreceptor lineage-specific transcription factor Otx2, thereby exhibiting neurogenic competence. At the molecular level, drug-induced dual activation of YAP and β-catenin elevated Cyclin D1 expression and repressed the cell cycle inhibitor p27^Kip1^, an effect mediated at least partially through the upregulation of the core transcriptional amplifier MYC. Collectively, this study established an efficient chemical strategy to boost adult mouse MG proliferation, offering a promising translational alternative for MG-mediated retinal regeneration.

## MATERIALS AND METHODS

### Animals

Specific-pathogen-free C57BL/6J mice and Glast-Cre^ERT2^;ROSA26R-tdTomato double-transgenic mice aged 6–8 weeks were used. Male and female mice were randomly included without sex stratification. Animals were housed under a 12-h light/12-h dark cycle with food and water ad libitum. All animals were housed in Xiamen University Laboratory Animal Center; all procedures were approved by the Xiamen University Institutional Animal Care and Use Committee (IACUC) (Approval No. XMULAC20230070) and complied with the *ARVO Statement for the Use of Animals in Ophthalmic and Vision Research.* Retinal explants were randomly assigned to treatment groups; exclusion criteria are described under Image acquisition and quantification.

### Genotyping

A 1–2⍰mm tail⍰tip or toe biopsy sample was lysed for PCR⍰based genotyping of the *Glast*⍰*CreE^RT2^* and *ROSA26R*⍰*tdTomato* alleles. Each 20 μL reaction system consisted of 10 μL 2× M⍰PCR OPTI Mix (Cat. No. B40013, Selleck, China), 0.5 μL of each of the four primers, 1 μL DNA template, and 7 μL ddH₂O. The thermal cycling profile was as follows: 94 °C for 5 min; nine touchdown cycles (94 °C for 30 s, annealing at 65 °C for 30 s with a 0.5 °C decrement per cycle, and 68 °C for 30 s); 27 cycles of 94 °C for 30 s, 60 °C for 30 s, and 72 °C for 30 s; followed by a final extension at 72 °C for 5 min and a hold at 4 °C. Amplicon sizes were approximately 600 bp for the Glast⍰CreERT2 transgene and 200 bp for the internal⍰control fragment; for ROSA26R⍰tdTomato, the wild⍰type and mutant amplicons were approximately 297 bp and 200 bp, respectively. Primer sequences are provided in **Supplementary Table** 3.

### Tamoxifen-induced MG lineage tracing

Tamoxifen was purchased from MedChemExpress (Cat. No. HY⍰13757A, China) and dissolved in corn oil to a concentration of 20 mg/mL at 37 °C under light⍰protected conditions. Glast⍰Cre^ERT2^;ROSA26R⍰tdTomato mice were given daily intraperitoneal injections of tamoxifen at a dose of 75 mg/kg for 5 consecutive days. Retinas were harvested one day following the final injection. All tdTomato lineage⍰tracing immunofluorescence assays were performed subsequent to this *in vivo* induction.

### Mouse retinal explant culture

Mice were deeply anesthetized using 2,2,2⍰tribromoethanol (Cat No. T48402, Sigma-Merck, Shanghai, China) and euthanized via cervical dislocation. Eyes were rapidly enucleated and transferred into HBSS (Cat No. 14025092, Thermo-Fisher, USA) or DMEM/F12 (Cat No. C11330500BT, Thermo-Fisher, China) supplemented with 1% penicillin⍰streptomycin (Cat No.15070063, Thermo-Fisher, USA). Neural retinas were isolated under aseptic conditions and radially cut at four sites to form a clover shaped and flattened explant. The time interval from tissue dissection to incubation was maintained within 1 h. Retinal explants were positioned ganglion cell layer⍰up and photoreceptor⍰layer⍰down onto sterile Falcon 40⍰μm cell strainers (Cat No.352340, Falcon, USA) in six⍰well plates. Each well contained 3 mL of culture medium consisting of Neurobasal⍰A (Cat No. 10888022, Lot No.3202812, Thermo-Fisher, USA), 2% B27 (Cat No. 17504044, Thermo-Fisher, USA), 1% N2 (Cat No. 17502048, Thermo-Fisher, USA), 2 mM GlutaMAX (Cat No. 35050061, Thermo-Fisher, USA), and 1% penicillin⍰streptomycin, and explants were cultured at 37 °C under 5% CO₂. Approximately half of the culture medium was replaced daily, with fresh drugs added with each medium change. Unless otherwise specified, one intact retina was defined as one experimental unit.

### Drug treatment and EdU labeling

TRULI (Cat. No. HY⍰138489, Lot No. 418329, MedChemExpress, China), CHIR⍰99021 (Cat. No. HY⍰10182, Lot No. 157150, MedChemExpress, China), Verteporfin (Cat. No. HY-B0146, MedChemExpress, China), ICG⍰001 (Cat. No. HY-14428, MedChemExpress, China), MYCi975 (Cat. No. SJ-MX0496, SparkJade, China), and EdU (Cat. No. B8337, APExbio, USA) were used in this study. All stock solutions were prepared in DMSO according to the manufacturer’s instructions and stored at −80 °C. Stock solutions were diluted with retinal explant culture medium to the following working concentrations: 10 μM TRULI, 6 μM CHIR⍰99021, 5 μM Verteporfin, 8 μM ICG⍰001, 5 μM MYCi975, and 10 μM EdU. For the control group, an equal volume of vehicle (DMSO, ∼0.1%) was added to the culture medium. All compounds were added on culture day 0 and replenished with each medium exchange. Treatment and sampling timelines are described in the figure legends.

### Retinal flat-mount immunofluorescence staining

Explants were fixed in 4% paraformaldehyde at room temperature for 2 h, rinsed with 1 PBS, and then blocked and permeabilized in 1 PBS containing 3% BSA and 0.5% Triton X⍰100 overnight at 4 °C. EdU Click⍰iT reactions were carried out using the Click⍰iT EdU Cell Proliferation Kits for Imaging with Alexa Fluor 488 (Cat. No. C10337, Thermo-Fisher/Invitrogen, USA) or Alexa Fluor 647 (Cat. No. C10340, Thermo-Fisher/Invitrogen, USA) for 2–3 h at room temperature under light⍰protected conditions. For immunofluorescence antibody staining, samples were incubated with primary antibodies overnight at 4 °C, followed by incubation with matching secondary antibodies (1:300 dilution) and Hoechst nuclear counterstain (1:2,000 dilution) for 2–3 h at room temperature in the dark. Retinal explants were flat⍰mounted along four quadrants and mounted in 50% glycerol for imaging acquisition. Retinal flat mounts were oriented with the photoreceptor layer facing the objective during confocal imaging.

### Cryo-sectioning and immunofluorescence

Retinas were fixed in 4% paraformaldehyde at room temperature for 2 h, followed by sequential cryoprotection in 5%, 10%, 20%, and 40% sucrose. Tissues were embedded in OCT compound (Cat No.3801480, lot No.010825, LeicaBiosystems, USA) and cryosectioned at 8 μm thickness. Sections were blocked in 1 PBS containing 3% BSA and 0.5% Triton X⍰100 for 1 h at room temperature. When required, a 30⍰min EdU click⍰iT reaction was carried out prior to antibody staining. Sections were incubated with primary antibodies overnight at 4 °C, then incubated with fluorophore⍰conjugated secondary antibodies (1:300 dilution) and Hoechst (1:2,000 dilution) for 1 h at room temperature in the dark. Sections in the central retina with the optic nerve head were preferentially selected for analysis.

### Western blotting

Each explant was lysed in approximately 60 μL RIPA buffer (50 mM Tris-HCl pH 7.5, 1 mM EDTA, 1 mM EGTA, 1% Triton X-100, 150 mM NaCl, 0.1% SDS, 1% sodium deoxycholate) supplemented with protease (Cat No. K1007, Lot No. BRN, APExbio, USA) and phosphatase inhibitor cocktails (Cat No.K1012, Lot No.26001K1012, APExbio, USA), homogenized on ice, and centrifuged at 14,000 rpm for 15 min at 4 °C. Protein concentrations were measured using the BCA Protein Assay Kit (Cat. No. 23227, Lot No. 3217963, Thermo Fisher/Pierce, USA). A total of 30 μg of protein was loaded per sample and resolved by 10% or 12% SDS⍰PAGE. PVDF membranes (Cat. No. 03010040001, Lot No. 73493400, Roche, UK) were activated in methanol for 2 min, equilibrated in transfer buffer, and subjected to wet⍰transfer at 90 V for approximately 120 min on ice. Following transfer, membranes were blocked in 5% non⍰fat milk for 2 h at room temperature with gentle rocking. Primary antibodies were diluted in 5% BSA, and membranes were incubated overnight at 4 °C. Membranes were washed three times for 10 min each in 1× TBST, incubated with HRP⍰tuconjugated secondary antibodies (1:5 000 dilution) for 2 h at room temperature, and washed three additional times in 1× TBST. Chemiluminescent signals were developed with the NcmECL SuperUltra Kit (Cat. No. P10060, NCM Biotech, China) and captured using the ChemiDoc Touch Imager (Bio Rad Laboratories Inc., Hercules, CA, USA). All antibodies used for Western blotting are listed in Supplementary Table 1. Band intensities were quantified with ImageJ (version 1.54f; National Institutes of Health, Bethesda, MD, USA) and normalized to loading controls GAPDH or α⍰Tubulin. Both normalized p YAP abundance and the p⍰YAP/YAP ratio were reported. Each Western blot comparison included a minimum of three independent retinal explants per group; exact n values are provided in the figure legends. Full⍰length, uncropped blot exposures are retained in the original source files and compiled in the Supplementary Figure 8 and 9.

### RNA extraction, reverse transcription, and qRT-PCR

Each biological replicate consisted of an individual retinal sample. Total RNA was isolated using Trizol Total RNA Extraction Reagent (Cat. No. RK30129, Abclonal, China). Subsequently, 1 μg of total RNA was reverse-transcribed into cDNA using the Hiaff® AdvanceFast 1st Strand cDNA Synthesis SuperMix for qPCR (DNA digester plus) (Cat. No. 11155ES60, Yeasen, China). Quantitative real-time PCR (qRT-PCR) was performed using Hieff^®^ qPCR SYBR Green Master Mix (No Rox) (Cat. No. 11201ES08, Yeasen, China) on a Roche LightCycler 96 system, with three technical replicates for each biological sample. The thermal cycling protocol was set as follows: an initial denaturation step at 95 °C for 30 s, followed by 40 cycles of 95 °C for 10 s, 60 °C for 20 s, and 72 °C for 20 s. A melting curve analysis was performed after amplification to verify PCR specificity. *Gapdh* was used as the internal reference gene, and relative gene expression levels were quantified via the 2^-△△Ct^ method. All samples shown in an individual figure were processed in a single batch, with no fewer than three independent retinal samples (n ≥ 3) per group. All primer sequences utilized in this study are listed in Supplementary Table 2.

### TUNEL Assay

TUNEL staining was performed on retinal cryosections using the Click-iT™ Plus TUNEL Assay for In Situ Apoptosis Detection with Alexa Fluor™ 647 dye (Cat. No. C10619, Invitrogen, Thermo Fisher Scientific, USA) according to the manufacturer’s instructions with minor modifications. Briefly, sections from pre-fixed retinal tissues were permeabilized with Proteinase K for 15 min, followed by TdT labeling for 60 min at 37 °C and Click-iT™ Plus detection for 30 min at 37 °C in the dark. Nuclei were counterstained with Hoechst before fluorescence imaging.

### RNA sequencing and bioinformatic analysis

Two independent retinas were included in each of the Ctrl, TRULI, CHIR, and TRULI+CHIR groups after 3 days of culture. RNA purification, library preparation, and sequencing were conducted by Majorbio Biotechnology Company, Shanghai, China. RNA-seq data have been deposited in the NCBI Sequence Read Archive (SRA) under BioProject accession PRJNA1521908. To identify differentially expressed genes (DEGs) between each group, the expression level of each transcript was calculated according to the transcripts per million reads (TPM) method. DEGs with |log2FC| ≥ 1.2 and *p*-adjust < 0.05 were considered to be significantly different expressed genes for the comparisons. In addition, functional enrichment analysis, including KEGG and GSEA, was performed based on DEGs, which were performed on the online platform of Majorbio Cloud Platform (https://cloud.majorbio.com/).

### Image acquisition and quantification

Fluorescence images were captured using an Olympus FV1000MPE upright multiphoton laser scanning confocal microscope (OLYMPUS Inc., Japan) equipped with a 20×/0.75 NA objective lens. Laser power, gain, offset, and scanning parameters were kept consistent across all experimental groups within each independent experiment. For retinal flat mount preparations, one imaging field was selected from each of the four retinal quadrants at the midpoint between the retinal periphery and the optic nerve head, generating four imaging fields per retina. For retinal cryosections, two fields were acquired on either side of the optic nerve head, also yielding four fields per retina. Cell counting and fluorescence intensity quantification were performed using ImageJ software. Regions exhibiting obvious tissue folds, damage, defocusing, or poor staining were excluded prior to unblinding. Cell counts from flat mount samples are presented as cells per square millimeter (cells/mm²), while counts from retinal sections are reported as cells per millimeter of retinal length (cells/mm). All image analyses were conducted by investigators blinded to the experimental group assignments.

For quantification of EdU+ tdTomato+ cells, the cell was classified as double-positive only when an EdU-labeled nucleus could be unambiguously assigned to a tdTomato-positive cell. Cells for which the EdU and tdTomato signals could not be confidently assigned to the same cell were not counted. Counts from the four fields were pooled to obtain one value per retina.

For quantitative analysis of Otx2 immunofluorescence signals, four independent retinas were analyzed per group at 7 days post-treatment. Individual regions of tdTomato^+^ MG in the Ctrl group and EdU^+^tdTomato^+^ MG in experimental groups were randomly selected, with approximately 100 cells quantified per group; each data point in the corresponding figure represents an individual cell. The mean Otx2 fluorescence intensity in tdTomato+ cells from the Ctrl group was defined as the normalization baseline, and the raw fluorescence intensity of each measured cell was normalized to this reference value.

### Statistical analysis

All statistical analyses excluding RNA⍰seq data were performed using GraphPad Prism 8.0 (GraphPad Software Inc., San Diego, CA, USA). All quantitative data are presented as the mean ± standard deviation (S.D.), as specified in the figure legends. Comparisons among multiple groups were performed using one⍰way ANOVA followed by Tukey’s multiple comparisons test, whereas two⍰group comparisons were conducted using a two⍰sided unpaired *t⍰*test. Statistical significance was defined as \**p* < 0.05, \*\**p* < 0.01, \*\*\**p* < 0.001, and \*\*\*\**p* < 0.0001; “ns” indicates no significant difference. Unless otherwise specified, each data point corresponds to one independent retinal sample.

## RESULTS

### TRULI and CHIR efficiently drive robust MG proliferation

To investigate the regulatory effects of Hippo inhibition and Wnt activation on MG proliferation and trans-differentiation, we examined the retinal expression of YAP and β⍰catenin, the core downstream transcriptional co-activators of the Hippo and Wnt signaling pathways, respectively. Immunostaining confirmed that both YAP and β⍰catenin were endogenously expressed in MG of adult murine retinas (**Figure S**1**A** and **B**).

We first applied TRULI (also known as Lats⍰IN⍰1), a selective Hippo pathway Lats kinase inhibitor that activates YAP and drives YAP-dependent MG proliferation in retinal organoids[18]. TRULI treatment reduced YAP phosphorylation in a dose-dependent manner (**Figure S**1**C**–**E**) and consequently upregulated canonical YAP target genes in cultured retinal explants (**Figure S**2**A**). Notably, TRULI alone exerted no significant influence on the expression of Wnt downstream targets (**Figure S**2**A**).

We next introduced CHIR⍰99021 (hereafter referred to as CHIR), a potent GSK3α/β inhibitor that stabilizes intracellular β⍰catenin and triggers Wnt pathway activation. CHIR treatment effectively elevated β⍰catenin protein levels in a dose⍰dependent manner (**Figure S**1**C** and **F**) and robustly induced the expression of Wnt target genes (**Figure S**2**B**). Intriguingly, CHIR alone significantly repressed the expression of YAP target genes including *Bmp4* and *Ctgf* (**Figure S**2**B**), indicating potential crosstalk and reciprocal regulation between the Hippo and Wnt pathways in the adult mouse retinas.

To specifically trace MG fate, we utilized the transgenic Glast⍰Cre^ERT2^; ROSA26R⍰tdTomato mouse line. Retinal explants were isolated from 6∼8-week-old transgenic mice and treated with vehicle control (DMSO), TRULI, CHIR, or the combined TRULI/CHIR regimen. The optimal combination of 10 μM TRULI and 6 μM CHIR was determined based on comprehensive evaluation of MG proliferative capacity and retinal cellular viability following 5 days of treatment (data not shown). After continuous 5-day EdU labeling, only very sparse EdU^+^tdTomato^+^ proliferative MG were detected in the DMSO control group (**Figure** 1**A**-**C**). While individual treatment with either TRULI or CHIR moderately promoted MG proliferation, combined TRULI and CHIR treatment elicited a robust increase in proliferative MG across the whole retina (**Figure** 1**C**). Quantitative analysis revealed that the control group exhibited approximately 5 EdU^+^tdTomato^+^ cells per mm^2^ in the central retina, whereas single⍰drug TRULI or CHIR treatment increased this density to ∼78 cells/mm² or ∼48 cells/mm², respectively. Strikingly, TRULI and CHIR combined treatment yielded a more than 15-fold increase in proliferative MG, reaching ∼1351 cells/mm² in the central retina (**Figure** 1**A** and **B**), indicating strong synergistic pro proliferative effects of TRULI and CHIR on adult retinal MG.

**Figure 1.**
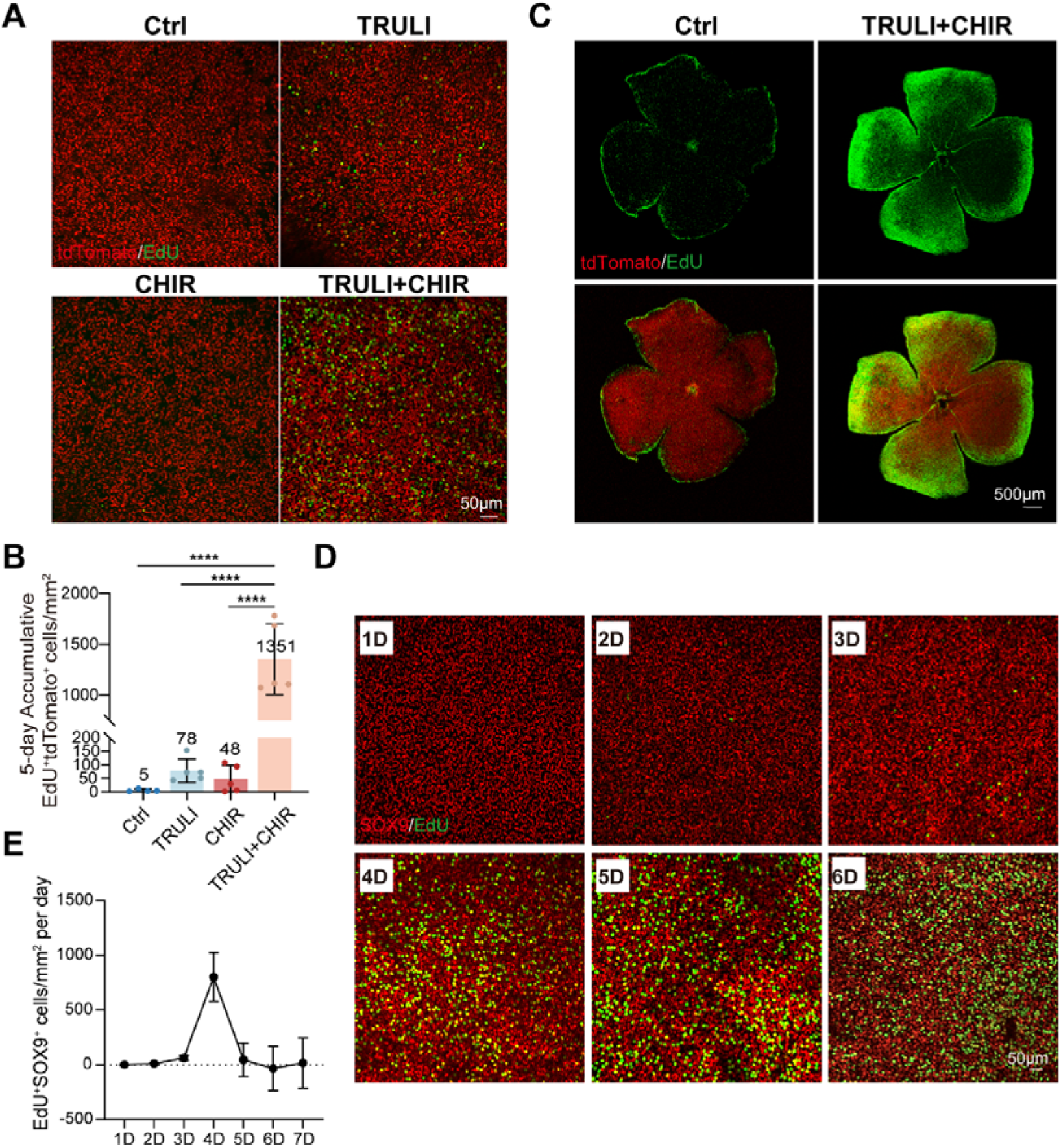
TRULI and CHIR synergistically promote robust Müller glia (MG) proliferation in murine retinal explants. (**A**) Analysis of EdU (*green*) incorporation in the retinal explants of ∼8-week Glast-Cre^ERT2^; tdTomato (*red*) mice after 5 days of Ctrl, TRULI, CHIR, or TRULI+CHIR treatment. (**B**) Quantification of the numbers of EdU⁺tdTomato⁺ cells per mm^2^ in the central retina in **A**. (**C**) Representative confocal images of EdU incorporation in the retinal wholemount after 5 days of Ctrl or TRULI+CHIR treatment. (**D**) Representative images of EdU (*green*) incorporation in SOX9 (*red*)-positive cells after TRULI+CHIR treatment with continuous EdU labeling for 1–6 days. (**E**) A time course analysis of MG proliferation by scoring the number of EdU+ SOX9^+^cells/mm^2^ on retinal flatmount following TRULI+CHIR treatment. In **B**, n=4 retinas for Ctrl and n=5 for each other group; in **E**, n=6 retinas per time point. Data are presented as mean ± SD. One-way ANOVA with Tukey’s multiple-comparisons test; \*\*\*\**p*<0.0001. Scale bars: 50 μm in **A** and **D**; 500 μm in **C**.

To characterize the kinetic profile of MG proliferation, retinal explants were subjected to continuous 7⍰day EdU incorporation assay under combined TRULI and CHIR treatment (**Figure** 1**D**). EdU^+^ proliferative cells emerged initially in the peripheral retina and gradually extended toward the central region over time (**Figure S**3**A**). Slightly different from previously reported *in vivo* observations[19], the density of EdU^+^ cells in our explant model peaked at day 4 post treatment (**Figure** 1**E**). Almost all EdU^+^ proliferative cells were immune-positive for Sox9, a canonical MG marker (**Figure** 1**D**), confirming that the proliferating population predominantly composed by retinal MG. To further validate the MG identity of proliferative cells, retinal explants were cryo-sectioned at a thickness of 8 μm. Consistently, nuclear EdU^+^ signals extensively colocalized with tdTomato^+^ labeling on retinal sections, further verifying the pro⍰proliferative effects of TRULI and CHIR on MG in adult mouse retinas (**Figure** 2**A** and **B**).

**Figure 2.**
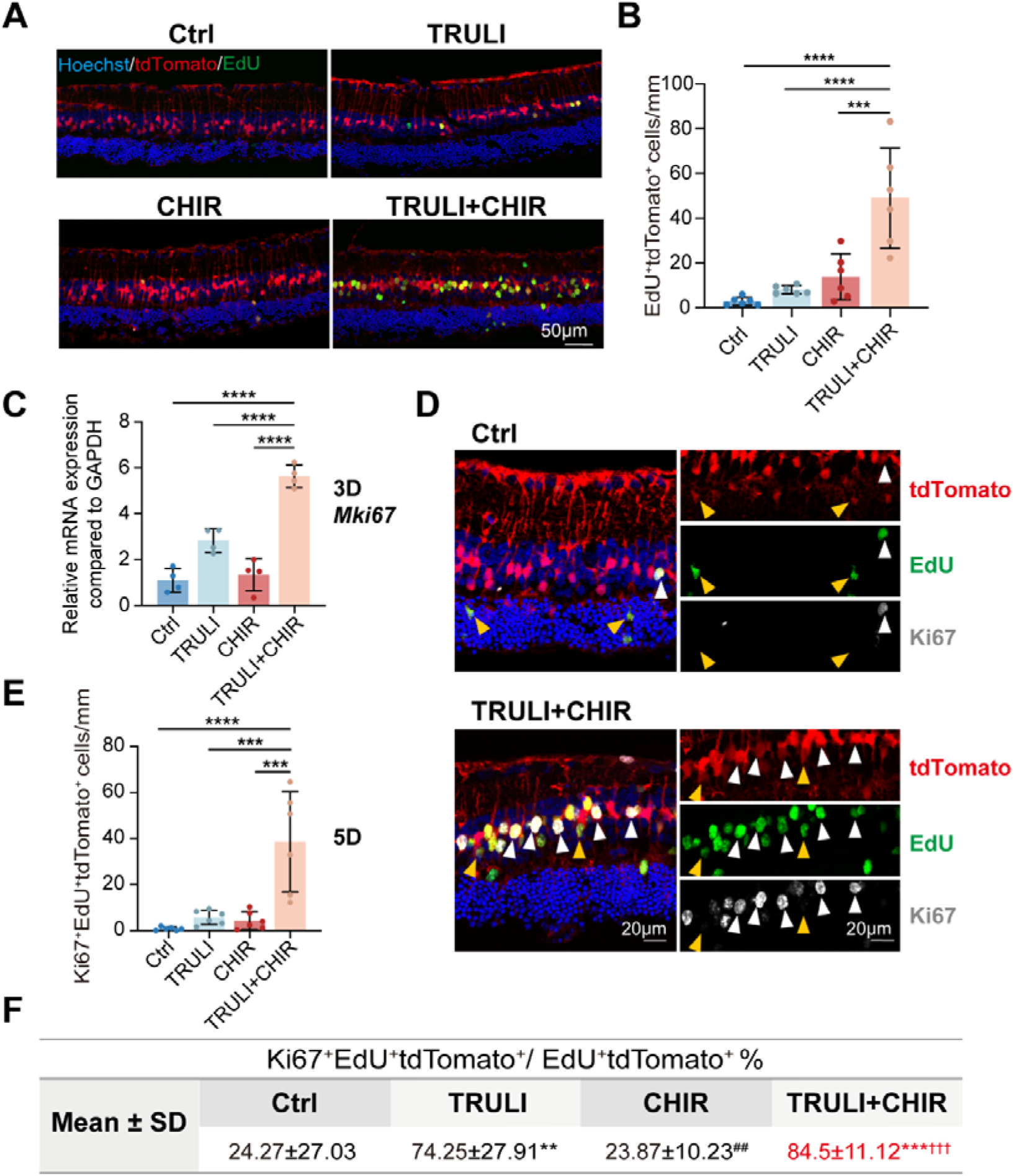
TRULI and CHIR robustly drive MG into the active cell cycle. (**A**) EdU (*green*) incorporation in Glast-Cre^ERT2^; tdTomato (*red*) labelled MG in retinal cross-sections after 5 days Ctrl, TRULI, CHIR, or TRULI+CHIR treatment. Cell nuclei were stained with Hoechst (*blue*). (**B**) Quantification of the numbers of EdU⁺tdTomato⁺ cells per mm in retinal sections in **A**. (**C**) qRT-PCR analysis of the mRNA levels of Mki67 after 3 days Ctrl, TRULI, CHIR, or TRULI+CHIR treatment. (**D**) Ki67 (*gray*) immunofluorescence and EdU (*green*) incorporation in Glast-Cre^ERT2^; tdTomato (*red*) labelled MG in retinal sections after 5 days Ctrl or TRULI+CHIR treatment; white arrows indicate Ki67^+^EdU^+^tdTomato^+^ cells and yellow arrows indicate EdU⁺tdTomato⁺ cells that were Ki67 negative. (**E**) Quantification of the numbers of Ki67⁺EdU⁺tdTomato⁺ cells per mm in retinal sections in **D**. (**F**) Quantification of the percentage of Ki67⁺EdU⁺tdTomato⁺ cells in EdU⁺tdTomato⁺ cells after 5 days Ctrl, TRULI, CHIR, or TRULI+CHIR treatment. n≥5 retinas per group in **B**, n=4 in **C**, and n=6 in **E** and **F**. Data are presented as mean ± SD; one-way ANOVA with Tukey’s multiple-comparisons test. In **F**, * denotes comparison with Ctrl, ^#^ with TRULI, and ^†^ with CHIR. \*\**p*<0.01, \*\*\**p*<0.001, \*\*\*\**p*<0.0001, ^##^*p*<0.01, ^†††^*p*<0.001. Scale bars: 50 μm in **A**; 20 μm in **D**.

We additionally employed Ki67, a canonical cell cycle marker, to confirm MG cell cycle re⍰entry. Quantitative real time PCR (qRT⍰PCR) results demonstrated significantly elevated *Mki67* mRNA levels at day 3 after combined TRULI and CHIR treatment (**Figure** 2**C**). Immunofluorescence staining further confirmed colocalized Ki67 expression in EdU^+^tdTomato^+^ cells at day 5 post treatment (**Figure** 2**D**). Quantification analyses also revealed that TRULI and CHIR combined group exhibited the highest density of Ki67^+^EdU^+^tdTomato^+^ triple positive cells (**Figure** 2**E**). Approximately 85% of all EdU^+^tdTomato^+^ cells were Ki67-positive following combined treatment, and this proportion was significantly higher than that in the control, TRULI only, and CHIR only groups (**Figure** 2**F**). Collectively, these results demonstrate that combined Hippo inhibition and Wnt activation effectively drives robust cell cycle re entry and proliferation of adult mouse retinal MG.

### Proliferating MG express retinal progenitor cell marker PAX6

In zebrafish, MG proliferation is coupled with their reprogramming toward retinal progenitor cells during endogenous retinal repair[1]. Inspired by this regenerative paradigm, we further investigated whether TRULI/CHIR⍰induced proliferative MG acquire retinal progenitor⍰like properties. Retinal explants were harvested three days after DMSO, TRULI, CHIR, or TRULI and CHIR combined treatment and subjected to bulk RNA⍰seq analysis. Gene set enrichment analysis (GSEA) revealed that gene sets associated with stemness signatures and cortical radial glial progenitor identity were enriched in the upregulated genes of the TRULI and CHIR combined group relative to the control group (**Figure** 3**A** and **B**). Consistent with the transcriptomic profiling, qRT⍰PCR validation confirmed the upregulation of multiple retinal stem/progenitor marker genes following TRULI and CHIR treatment. Interestingly, different from zebrafish which upregulated Ascl1 after retinal damage, *Neurog2* exhibited the most prominent induction after TRULI and CHIR treatment in mouse retinal explants (**Figure S**3**B**). These collective findings indicate that proliferative MG undergo cellular reprogramming and acquire retinal progenitor cell (RPC)-like characteristics.

**Figure 3.**
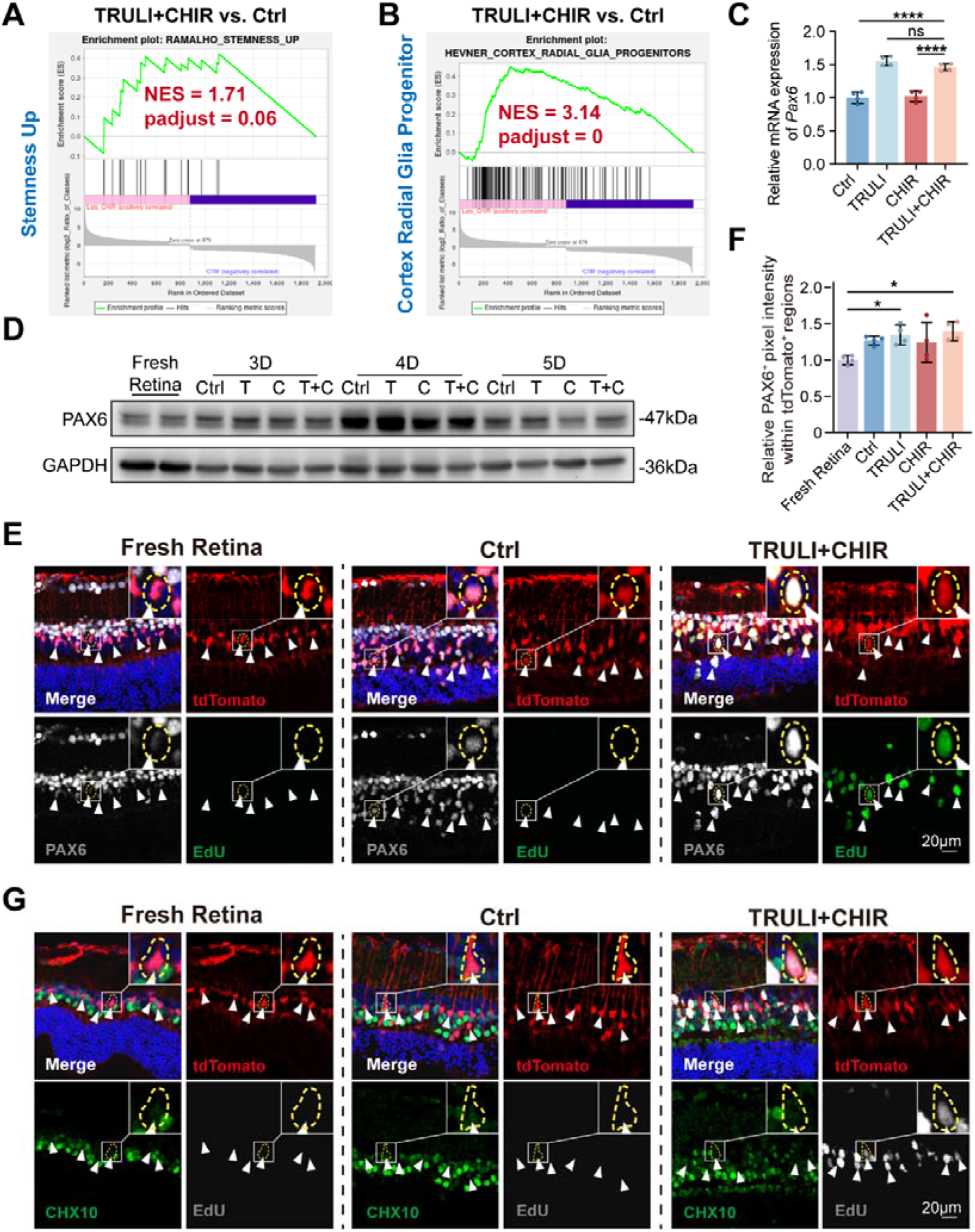
MG enter into progenitor-like state. (**A**,**B**) Gene-set enrichment analysis(GSEA) of RNA-seq samples after 3 days Ctrl or TRULI+CHIR treatment shows the enrichment of genes in Stemness Up and Cortex Radial Glia Progenitor, NES, normalized enrichment score; n=2 retinas per group. (**C**) qRT-PCR analysis of the mRNA levels of *Pax6* after 3 days Ctrl, TRULI, CHIR, or TRULI+CHIR treatment. (**D**) Western blot analysis of the protein levels of PAX6 in fresh retina and retinal explants after 3–5 days Ctrl, TRULI(T), CHIR(C), or TRULI+CHIR(T+C) treatment; GAPDH was used as loading control. (**E**) Immunofluorescence of PAX6 (*gray*) and EdU (*green*) incorporation in Glast-Cre^ERT2^; tdTomato (*red*) labelled MG in retinal sections of fresh retina or after 5 days Ctrl or TRULI+CHIR treatment. Cell nuclei were stained with Hoechst (*blue*). White arrows indicate PAX6^+^tdTomato^+^ cells. (**F**) Quantification of Relative PAX6 fluorescence intensity within tdTomato⁺ regions in **E**; n=4 retinas per group. (**G**) Immunofluorescence of CHX10 (*green*) and EdU (*gray*) incorporation in Glast-Cre^ERT2^; tdTomato (*red*) labelled MG in retinal sections of fresh retina or after 5 days Ctrl or TRULI+CHIR treatment. Cell nuclei were stained with Hoechst (*blue*); no clear CHX10 induction was detected in any group. Data are presented as mean ± SD; one-way ANOVA with Tukey’s multiple-comparisons test. \**p*<0.05, \*\*\*\**p*<0.0001; ns, not significant. Scale bars: 20 μm in **E** and **G**.

PAX6 serves as a core master transcription factor governing ocular development, maintaining the multipotent state of RPCs and orchestrating retinal neurogenesis throughout development[20, 21]. On the transcriptional level, both TRULI only or TRULI and CHIR combined treatment significantly upregulated *Pax6* mRNA expression at day 3 post treatment (**Figure** 3**C**). Western blotting further detected a sharp increase in pan⍰retinal PAX6 protein levels at day 4 across all experimental groups, a time point coinciding with robust MG cell cycle re-entry (**Figure** 3**D**). However, there was no significant difference in global PAX6 protein abundance between the TRULI and CHIR combinatorial treatment and control groups at this time point (**Figure** 3**D**). At day 5 post treatment, retinal cryo-sections were examined to characterize the spatial distribution of PAX6 expression. Quantitative analysis verified that PAX6 expression was specifically and sustained elevated in tdTomato^+^ MG following TRULI and CHIR treatment (**Figure** 3**E** and **F**). In contrast, another canonical RPC marker CHX10 was undetectable in the activated MG under the same treatment conditions (**Figure** 3**G**).

In the mammalian retina, retinal injury typically triggers reactive gliosis in resident MG, which is characterized by rapid upregulation of the intermediate filament protein GFAP [2]. Despite robust cell cycle re⍰entry in TRULI-and-CHIR⍰activated MG, qRT⍰PCR results revealed significant suppression of GFAP expression upon combined TRULI and CHIR treatment (**Figure S**3**C**-**E**). The repression of gliotic signatures further supports the claim that TRULI and CHIR stimulation drives MG reprogramming toward an RPC like state rather than initiating reactive gliosis.

### TRULI-and-CHIR-treated MG show modest upregulation of Otx2, a transcription factor specific to the bipolar and photoreceptor lineages

In zebrafish, proliferating MG transiently acquire an RPC-like state, exit the cell cycle, and ultimately undergo neuronal trans-differentiation to restore retinal function[1]. To determine whether TRULI-and-CHIR-activated MG possess neurogenic potential, we further traced their cell fate during culture. Following 5 days of continuous TRULI and CHIR treatment, robust EdU nuclear incorporation was detected in a large population of MG. Nevertheless, nearly all EdU^+^tdTomato^+^ cells remained positive for the canonical MG marker Sox9, indicating that these proliferative cells largely retained glial identity at this early stage (**Figure S**3**F**).

We next performed immunofluorescence staining for multiple neuronal markers to evaluate early trans-differentiation events, including the pan-neuronal marker NeuN, the bipolar/photoreceptor lineage marker Otx2, and the amacrine/ganglion cell marker HuC/D. At day 5 post treatment, only a small subset of EdU-positive MG exhibited weak Otx2 expression, whereas no detectable NeuN or HuC/D signals were observed in these proliferative cells (**Figure** 4**A**, **Figure S**4**A** and **B**).

**Figure 4.**
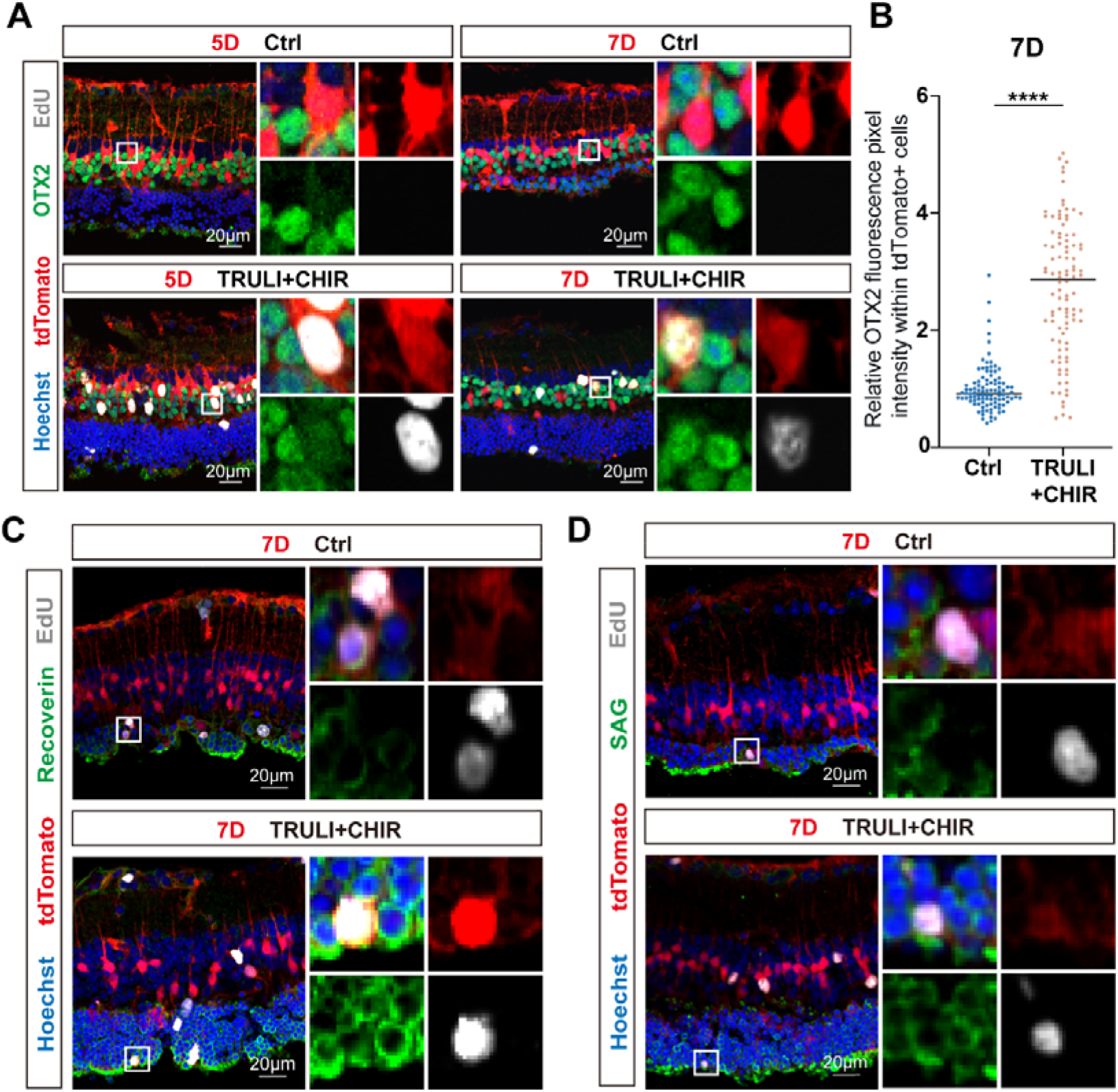
Trans-differentiation of proliferating MG into neuron-like cells. (**A**) Immunofluorescence of OTX2 (*green*) and EdU (*gray*) incorporation in Glast-Cre^ERT2^; tdTomato (*red*) labelled MG in retinal sections after 5 days CTRL or TRULI+CHIR treatment as well as follow-up to day 7. Cell nuclei were stained with Hoechst (*blue*). (B) Quantification of relative OTX2 fluorescence intensity in tdTomato⁺ cells on day 7. Measured cells were obtained from four independent retinas per group, with approximately 100 cells analyzed per group; each point represents one cell. (**C**) Immunofluorescence of Recoverin (*green*) and EdU (*gray*) incorporation in Glast-Cre^ERT2^; tdTomato (*red*) labelled MG in retinal sections on day 7. Cell nuclei were stained with Hoechst (*blue*). (**D**) Immunofluorescence of SAG (*green*) and EdU (*gray*) incorporation in Glast-Cre^ERT2^; tdTomato (*red*) labelled MG in retinal sections on day 7. Cell nuclei were stained with Hoechst (*blue*). Student’s *t*-test. \*\*\*\**p*<0.0001. Scale bars: 20 μm.

Subsequently, retinal explants culture was extended for an additional 2 days after finishing the 5-day drug treatment. At day 7, Otx2 expression was markedly upregulated in EdU^+^tdTomato^+^ MG after combined TRULI and CHIR treatment (**Figure** 4**A**). Although sporadic weak Otx2 signals were present in control MG, quantitative analysis confirmed that TRULI and CHIR treatment significantly increased Otx2 expression levels in tdTomato^+^ MG (**Figure** 4**B**). In contrast, proliferative MG remained negative for HuC/D expression throughout the culture period, indicating no differentiation toward amacrine or ganglion cell lineages (**Figure S**4**B**). Notably, a few EdU^+^tdTomato^+^ cells in the combined TRULI and CHIR group expressed the mature photoreceptor markers Recoverin and SAG (**Figure** 4**C** and **D**), suggesting that TRULI-and-CHIR-activated MG may be capable of initiating trans-differentiation toward rod photoreceptor fate.

Furthermore, TUNEL staining was performed to exclude the possibility that altered cell fate resulted from selective MG cell death. TUNEL-positive apoptotic cells were predominantly localized in the outer nuclear layer (ONL) rather than in reprogrammed MG populations, indicating that the observed neurogenic potential towards bipolar and photoreceptor lineage but not amacrine and ganglion cell lineage of activated MG was not caused by MG depletion or cell apoptosis (**Figure S**5).

### TRULI and CHIR enhance YAP and β-catenin nuclear translocation and transcriptional activities to promote MG proliferation

To dissect the molecular mechanism by which TRULI and CHIR co-regulate MG cell cycle re-entry, we first characterized the activation status of their core downstream effectors, YAP and β-catenin. Time-course Western blot analysis demonstrated that TRULI treatment reduced YAP phosphorylation, while CHIR treatment effectively elevated total β-catenin protein levels (**Figure** 5**A** and **B**). In the TRULI and CHIR group, the reduction in phosphorylated YAP was comparable to that observed in the TRULI single-treatment group, whereas the upregulation of β-catenin was moderately attenuated relative to CHIR single-agent treatment (**Figure** 5**A** and **B**). These results suggest potential signaling crosstalk between YAP and β-catenin upon dual pathway activation.

**Figure 5.**
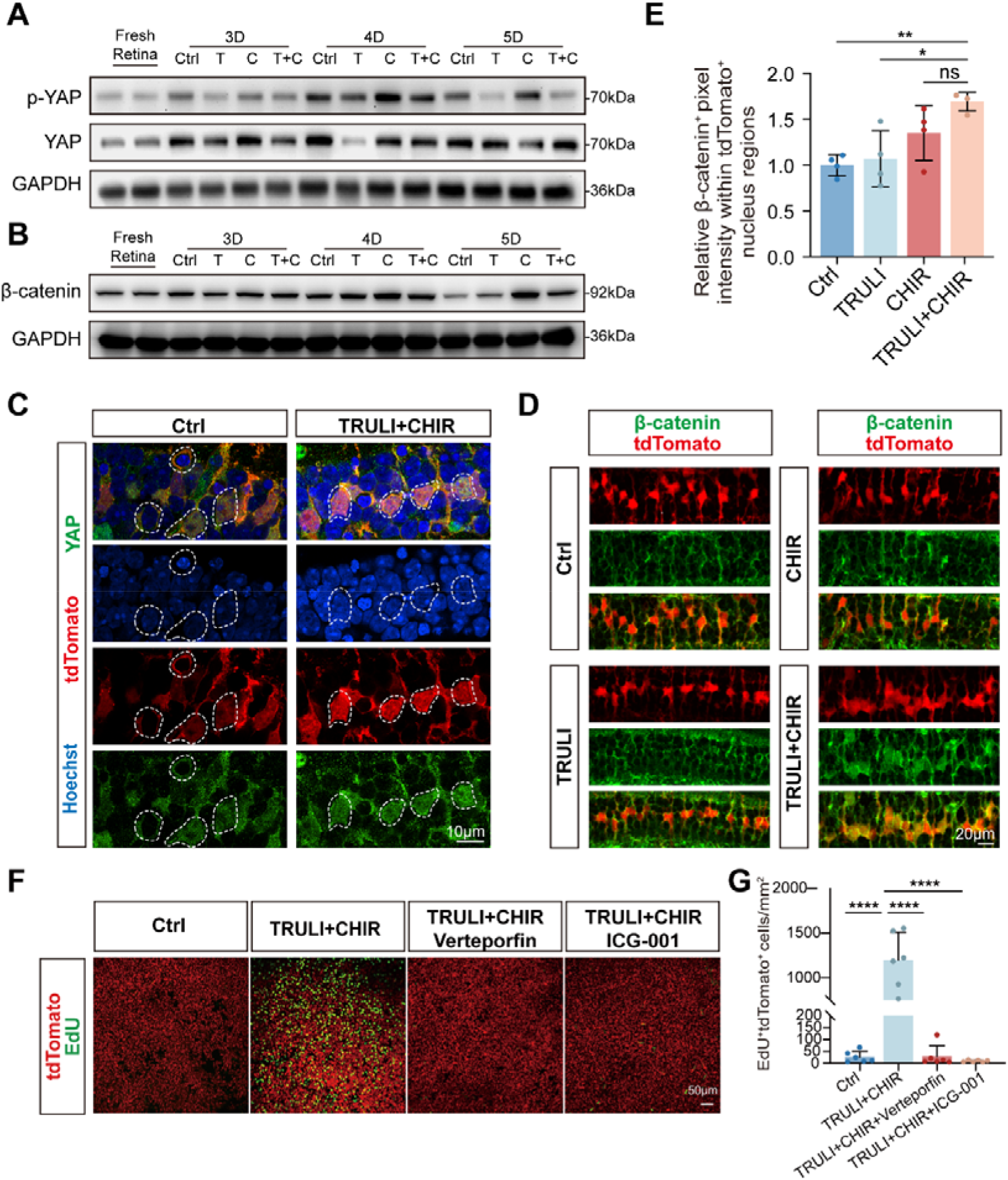
TRULI and CHIR treatment enhances YAP and β-catenin nuclear translocation and transcriptional activities. (**A**) Western blot analysis of p-YAP and YAP levels in fresh retina or retinal explants after 3–5 days Ctrl, TRULI(T), CHIR(C), or TRULI+CHIR(T+C) treatment. GAPDH was used as loading control. (**B**) Western blot analysis of β-catenin levels in fresh retina or retinal explants after 3–5 days Ctrl, TRULI(T), CHIR(C), or TRULI+CHIR(T+C) treatment. GAPDH was used as loading control. (**C**) Immunofluorescence of YAP (*green*) in Glast-Cre^ERT2^; tdTomato (*red*) labelled MG in retinal sections after 5 days CTRL or TRULI+CHIR treatment. Cell nuclei were stained with Hoechst (*blue*). (**D**) Immunofluorescence of β-catenin (*green*) in Glast-Cre^ERT2^; tdTomato (*red*) labelled MG in retinal sections after 5 days CTRL, TRULI, CHIR, or TRULI+CHIR treatment. (**E**) Relative β-catenin fluorescence intensity in tdTomato⁺/Hoechst⁺ overlapping nuclear regions; n=4 retinas per group. (**F**) EdU (*green*) incorporation in Glast-Cre^ERT2^; tdTomato (*red*) labelled MG in retinal flat wholemounts after 5 days of Ctrl, TRULI+CHIR, TRULI+CHIR with Verteporfin or TRULI+CHIR with ICG-001 treatment. (**G**) Quantification of the numbers of EdU⁺tdTomato⁺ cells per mm^2^ in the central retina in **F.** n=6 retinas per group. Data are presented as mean ± SD; one-way ANOVA with Tukey’s multiple-comparisons test. \**p*<0.05, \*\**p*<0.01, \*\*\*\**p*<0.0001; ns, not significant. Scale bars: 10 μm in **C**; 20 μm in **D**; 50 μm in **F**.

Consistent with reduced YAP phosphorylation, immunofluorescence staining revealed enhanced nuclear accumulation of YAP in tdTomato^+^ MG following combined TRULI and CHIR treatment (**Figure** 5**C**). Similarly, β-catenin exhibited a pronounced subcellular translocation: it was predominantly distributed in the cytoplasm and cell membrane under basal conditions, but accumulated prominently in the nucleus after combinatorial drug treatment (**Figure** 5**D** and **E**). Quantitative analysis of nuclear β-catenin signals in tdTomato^+^Hoechst^+^ nuclear regions of MG confirmed significantly higher nuclear β-catenin abundance in the TRULI and CHIR combinatorial treatment group compared with the control and TRULI only groups. Although the combinatorial group also showed a slight increase in nuclear β-catenin relative to CHIR only treatment, this difference did not reach statistical significance (**Figure** 5**E**).

Nuclear translocation of YAP and β-catenin is essential for their transcriptional co-activator functions. We further investigated whether YAP- and β-catenin-dependent transcriptional activation mediates TRULI-and-CHIR-induced MG proliferation. To this end, we applied two specific pathway inhibitors: Verteporfin, which disrupts the endogenous YAP–TEAD interaction to block YAP transcriptional activity [22], and ICG-001, which selectively suppresses β-catenin/TCF-mediated downstream transcription[23]. qRT-PCR analysis verified that 4 days of TRULI and CHIR treatment upregulated both canonical YAP target genes (*Bmp4*, *Cyr61*, *Ctgf*) and Wnt/β-catenin target genes (*Dkk1*, *Axin2*, *Lef1*) (**Figure S**6). Pharmacological inhibition with Verteporfin or ICG-001 markedly changed the expression of these target genes, confirming the specific regulatory effects of the two inhibitors on YAP and β-catenin transcriptional activity (**Figure S**6). Notably, Verteporfin and ICG-001 induced highly consistent patterns of target gene expression alterations, indicating that YAP and β-catenin may act cooperatively to control downstream transcriptional programs during TRULI-and-CHIR-induced MG activation. Consistently, blockage of either YAP-TEAD or β-catenin/TCF signaling almost completely abolished MG proliferation triggered by TRULI and CHIR treatment (**Figure** 5**F** and **G**). Collectively, these results demonstrate that YAP- and β-catenin-dependent transcriptional activation is indispensable for the pro-proliferative effects of dual Hippo/Wnt pathway modulation in adult retinal MG.

### The upregulation of Cyclin D1 and the downregulation of p27^Kip1^ underlie MG cell cycle reentry after TRULI and CHIR treatment

To systematically identify the core molecular drivers underlying TRULI-and-CHIR-induced MG proliferation, we performed transcriptomic profiling via RNA-seq and comparative analysis of differentially expressed genes (DEGs) between control and TRULI-and-CHIR-treated retinal explants. KEGG pathway enrichment analysis revealed significant enrichment of the cell cycle pathway (mmu04110, padjust = 0.0007) in DEGs derived from the combinatorial group (**Figure** 6**A**). Hierarchical clustering of cell cycle-related DEGs further demonstrated widespread transcriptional alterations in core cell cycle regulators, including cyclins, cyclin-dependent kinases, and cell cycle checkpoint inhibitors (**Figure** 6**B**). Among these genes, two well-characterized regulators of MG proliferation were prioritized: *Ccnd1* (encoding Cyclin D1) and *Cdkn1b* (encoding p27^Kip1^). Previous studies have established that YAP activation triggers Cyclin D1 upregulation to drive MG cell cycle re-entry in the adult mouse retina[12, 13], while concurrent p27^Kip1^ downregulation and cyclin activation are essential for sustained MG cell cycle progression[2, 14]. We therefore focused on these two genes to delineate the downstream cell cycle machinery modulated by the combined TRULI and CHIR treatment. Quantitative RT-PCR validation at day 3 confirmed that *Ccnd1* expression was significantly elevated in the TRULI and CHIR combinatorial treatment group compared with the control group, and this inductive effect was comparable to that observed in TRULI only treatment(**Figure** 6**C**), In contrast, CHIR only treatment did not alter the expression of *Ccnd1* gene (**Figure 6C**). In contrast, *Cdkn1b* mRNA levels exhibited no significant transcriptional changes across experimental groups (**Figure** 6**D**). Consistent with the transcriptional trend of cyclin genes, Western blot analysis at day 4 demonstrated Cyclin D1 protein upregulation and marked p27^Kip1^ protein repression following TRULI and CHIR treatment (**Figure** 6**E**–**H**).

**Figure 6.**
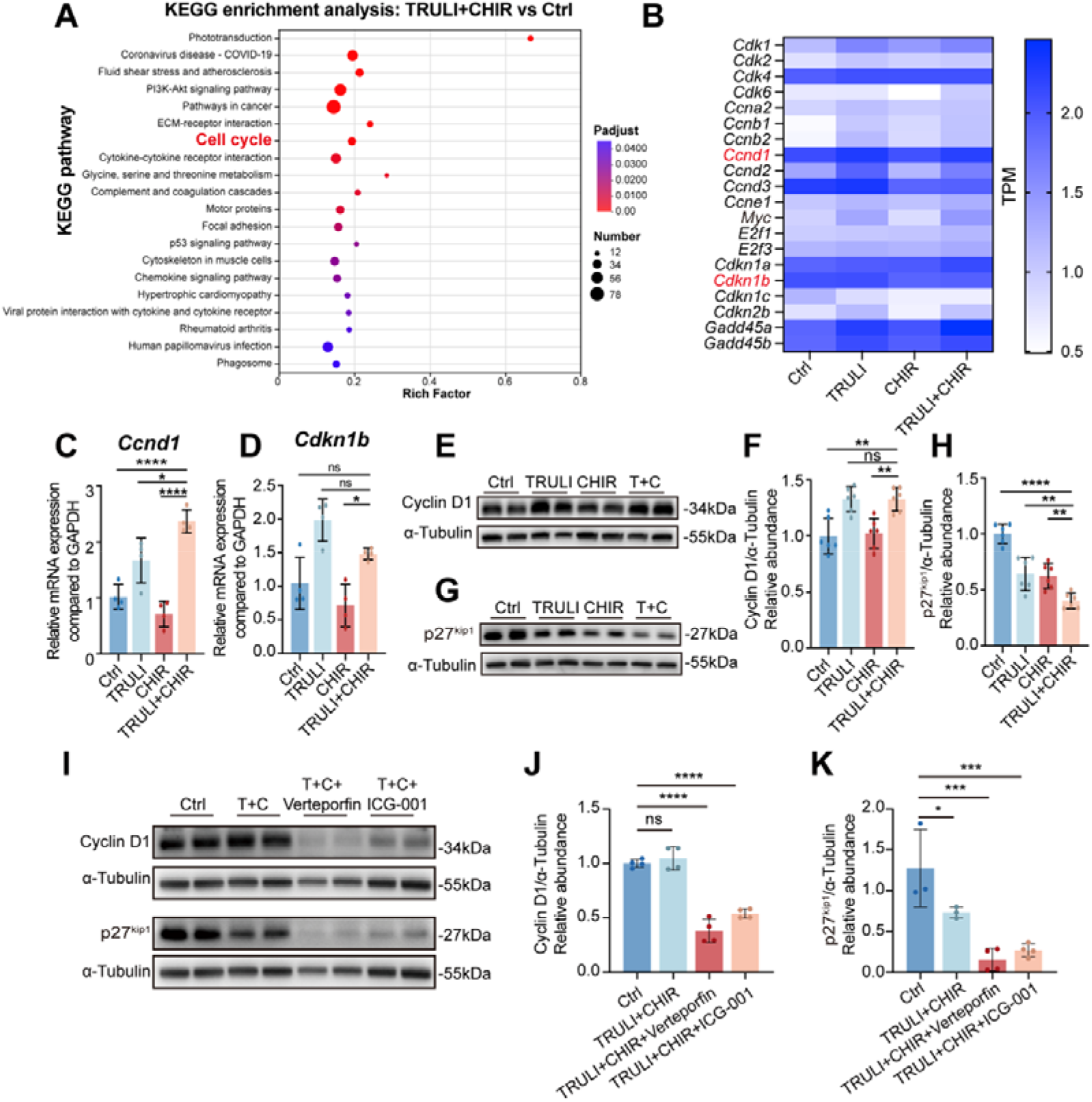
TRULI and CHIR treatment promotes MG cell-cycle reentry by modulating the expression levels of Cyclin D1 and p27^kip1^. (**A**) KEGG enrichment analysis of differentially expressed genes (DEGs) in TRULI+CHIR group relative to Ctrl group after 3 days treatment. n= 2 retinas per group. (**B**) Heat map of enriched differentially expressed cell-cycle-associated genes. (**C**, **D**) qRT-PCR analysis of mRNA levels of *Ccnd1* and *Cdkn1b* in the retinal explants after 3 days Ctrl, TRULI, CHIR, or TRULI+CHIR treatment. n=4 retinas per group. (**E**, **F**) Western blot analysis and quantification of protein levels of Cyclin D1 after 4 days Ctrl, TRULI, CHIR, or TRULI+CHIR (T+C) treatment; α-Tubulin was used as loading control. n=6 retinas per group. (**G**, **H**) Western blot analysis and quantification of protein levels of p27^kip1^ after 4 days Ctrl, TRULI, CHIR, or TRULI+CHIR (T+C) treatment; α-Tubulin was used as loading control. n=6 retinas per group. (**I**) Western blot analysis of Cyclin D1 and p27^kip1^ protein levels in retinal explants after 4 days Ctrl, TRULI+CHIR (T+C), TRULI+CHIR+Verteporfin (T+C+Verteporfin), or TRULI+CHIR+ICG-001 (T+C+ICG-001) treatment; α-Tubulin was used as loading control. (**J**, **K**) Quantification of relative Cyclin D1 and p27^kip1^ protein levels in **I**. n = 3 retinas for Ctrl and TRULI+CHIR group in **K**, and n=4 retinas for all other groups. Data are presented as mean ± SD; one-way ANOVA with Tukey’s multiple-comparisons test. \**p*<0.05, \*\**p*<0.01, \*\*\**p*<0.001, \*\*\*\**p*<0.0001; ns, not significant.

To determine whether YAP and β-catenin transcriptional activity governs the regulation of Cyclin D1 and p27^Kip1^, we examined the expression changes of these two cell cycle regulators upon pharmacological blockade of YAP-TEAD or β-catenin/TCF signaling. Inhibition of YAP or β-catenin transcriptional activity effectively abrogated TRULI-and-CHIR-induced Cyclin D1 upregulation at both the mRNA and protein levels (**Figure** 6**I** and **J**, **Figure S**6**G**). Intriguingly, pharmacological suppression of YAP/β-catenin-mediated transcription further reduced p27^Kip1^ expression at both transcriptional and translational levels (**Figure** 6**I** and **K**, **Figure S**6**H**), implying that additional transcription-independent mechanisms participate in the regulation of p27^Kip1^ under dual pathway activation.

### MYC serves as a central downstream node mediating MG proliferation after TRULI and CHIR treatment

KEGG pathway analysis of DEGs also identified enrichment of Hippo (mmu04390) and Wnt (mmu04310) signaling pathways in the TRULI and CHIR group, although the enrichment did not reach statistically significant level (Hippo: padjust = 0.10; Wnt: padjust = 0.56). Comparative analysis of overlapping DEGs between the two pathways revealed significant upregulation of *Myc*, a master transcriptional regulator governing cell proliferation (**Figure** 7**A** and **B**)[24]. Consistently, gene set enrichment analysis (GSEA) confirmed robust enrichment of MYC target gene signatures in the TRULI and CHIR combinatorial treatment group (NES = 2.64, padjust = 0), further supporting the pivotal role of MYC in driving MG proliferative reprogramming (**Figure** 7**C**). qRT-PCR analysis confirmed that *Myc* mRNA expression was significantly upregulated at day 3 following the combined TRULI and CHIR treatment, with substantially higher levels than those in single-treatment and control groups (**Figure** 7**D**). Western blot analysis further verified prominent induction of both C-MYC and N-MYC protein expression at day 4 post TRULI and CHIR treatment (**Figure** 7**E** and **F**). Consistently, immunofluorescence staining demonstrated specific and robust elevation of MYC protein signals in tdTomato^+^ MG after TRULI and CHIR treatment (**Figure** 7**G**). These results suggest MYC as a potential intermediate node that bridges upstream YAP/β-catenin transcriptional activation and downstream cell cycle remodeling.

**Figure 7.**
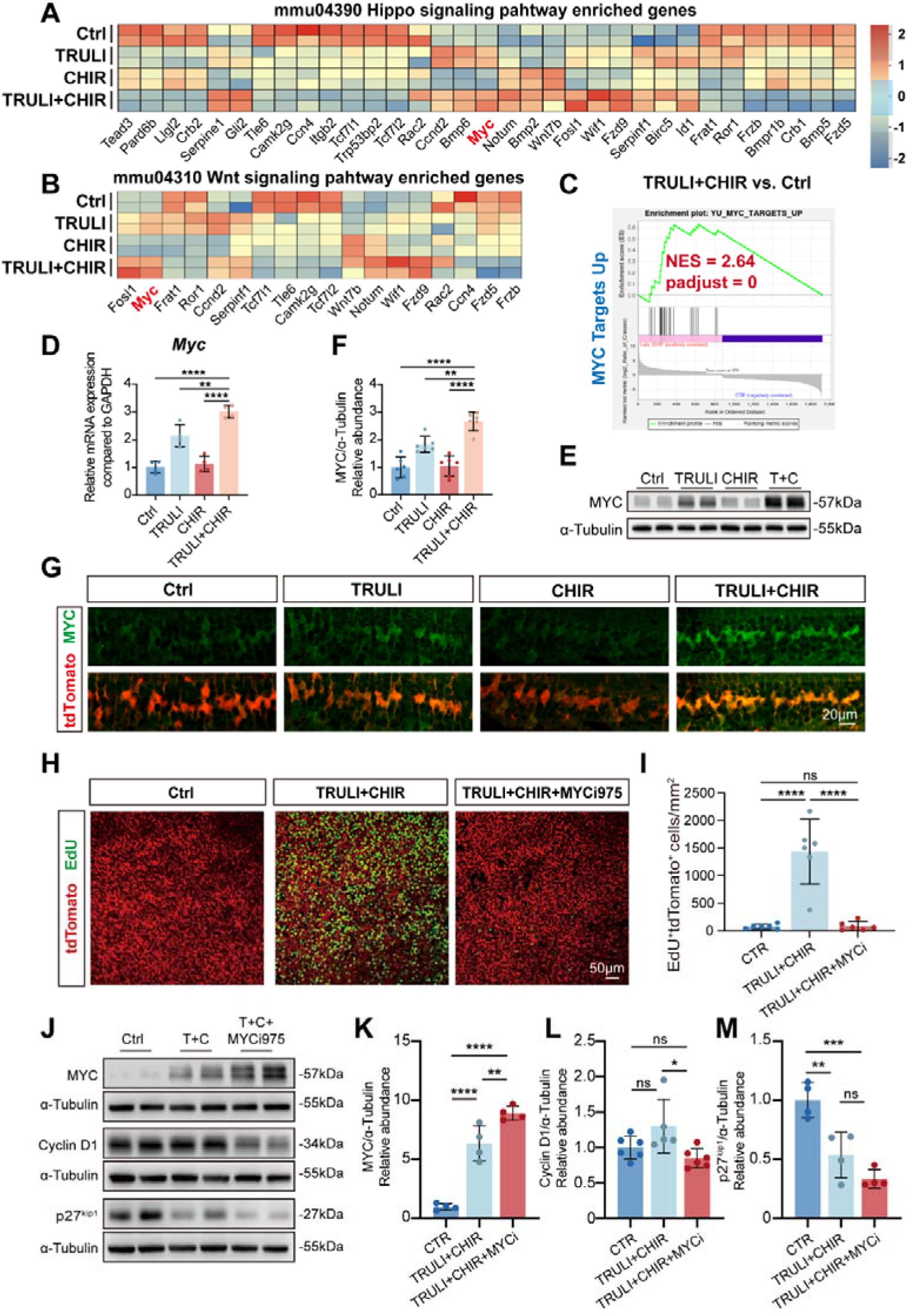
MYC is a key node underlying TRULI+CHIR-induced MG cell-cycle re-entry. (**A**,**B**) Heat maps of Hippo- and Wnt-associated differentially expressed genes. (**C**) GSEA analysis shows enrichment of MYC target genes (NES=2.64, adjusted *p*-value=0). (**D**) qRT-PCR analysis of mRNA levels of *Myc* in retinal explants after 3 days Ctrl, TRULI, CHIR, or TRULI+CHIR treatment. n= 4 retinas per group. (**E**) Western blot analysis of MYC protein levels in retinal explants after 4 days Ctrl, TRULI, CHIR, or TRULI+CHIR treatment; α-Tubulin was used as loading control. (**F**) Quantification of relative MYC protein levels in **E**. n= 6 retinas per group. (**G**) Immunofluorescence of MYC (*green*) in Glast-Cre^ERT2^; tdTomato (*red*) labelled MG in retinal sections after 5 days of Ctrl, TRULI, CHIR, or TRULI+CHIR treatment. (**H**) EdU (green) incorporation in Glast-Cre^ERT2^; tdTomato (*red*) labelled MG in retinal flat wholemounts after 5 days of Ctrl, TRULI+CHIR, or TRULI+CHIR with MYCi975 treatment. (**I**) Quantification of the numbers of EdU⁺tdTomato⁺ cells per mm^2^ in the central retina in **H.** n=6 retinas per group. (**J**) Western blot analysis of MYC, Cyclin D1, and p27^kip1^ protein levels in retinal explants after 4 days Ctrl, TRULI+CHIR, or TRULI+CHIR with MYCi975 treatment; α-Tubulin was used as loading control. (**K**,**L**,**M**) Quantification of relative MYC, Cyclin D1 and p27kip1 protein levels in **J**. n ≥ 4 retinas per group. Data are presented as mean ± SD; one-way ANOVA with Tukey’s multiple-comparisons test. \**p*<0.05, \*\**p*<0.01, \*\*\*\**p*<0.0001; ns, not significant. Scale bars: 20 μm in **G**; 50 μm in **H**

To functionally test the requirement of MYC for TRULI-and-CHIR-triggered MG proliferation, we used MYCi975, a selective small-molecule inhibitor that disrupts MYC/MAX dimerization and suppresses MYC-dependent transcriptional programs[25]. MYCi975 administration almost completely abolished TRULI-and-CHIR-induced EdU incorporation in MG (**Figure** 7**H**). Quantitative analysis showed that the density of EdU^+^tdTomato^+^ proliferative MG decreased sharply from approximately 1436 cells/mm² in the TRULI and CHIR combinatorial treatment group to 83 cells/mm² when MYCi975 was co-treated, a value comparable to the basal proliferative level of 77 cells/mm² in control retinas (**Figure** 7**I**). Interestingly, MYC inhibition led to a moderate increase in MYC mRNA and protein expression (**Figure** 7**J** and **K**, **Figure S**7**A**), which likely resulted from endogenous competition between MYC transcriptional suppression and YAP/β-catenin transcriptional activation. Importantly, MYCi975 treatment markedly downregulated Cyclin D1 protein levels and further repressed p27^Kip1^ expression, accompanied by abrogation of MG cell cycle re-entry (**Figure** 7**J**, **L** and **M**). Collectively, the phenotypic and molecular alterations induced by MYC inhibition closely recapitulated the inhibitory effects of YAP/β-catenin transcriptional blockade, demonstrating that MYC acts as an essential downstream effector to mediate the pro-proliferative and reprogramming activity of TRULI and CHIR in adult retinal MG.

## DISCUSSION

In zebrafish, retinal MG possess robust endogenous regenerative capacity, wherein injury-induced MG proliferation and reprogramming drive progenitor formation and neuronal differentiation in a sequential way to replenish lost retinal neurons. In mammals, however, MG regenerative potential is severely restricted, with insufficient cell cycle re-entry serving as a primary barrier for retinal repair [2]. Although multiple genetic strategies have been established to promote mammalian MG proliferation and reprogramming [12–14], most transgenic manipulations tend to lock MG in a persistent proliferative state and fail to support subsequent cell cycle exit and postmitotic neuronal differentiation. Here, we demonstrate that combined treatment with the Hippo signaling inhibitor TRULI and the Wnt signaling activator CHIR 99021 efficiently triggered robust proliferation in adult mouse MG. Compared with integrating genetic interventions, small-molecule-based chemical modulation offers distinct translational advantages, including non-invasiveness, chromosomal integrity, and functional reversibility[26]. While previous studies have reported that growth factors such as EGF, FGF, and FGF/insulin combinations can induce murine MG proliferation [11], synthetic small molecules exhibit superior controllability, standardizability, and cost-efficiency, making chemical cocktails a promising strategy for future clinical retinal regeneration applications.

MYC is a well-characterized core regulator of cell cycle progression, pluripotency maintenance, and somatic cell reprogramming, with essential roles in induced pluripotent stem cell generation [27]. Notably, MYC-mediated regenerative programs are evolutionarily conserved across vertebrates. In zebrafish, the mammalian MYC ortholog *Mycb* is significantly upregulated after retinal injury and functions as a critical upstream driver of MG proliferation and reprogramming[28, 29]. In mammals, three MYC paralogs (c-MYC, N-MYC, and L-MYC) exert distinct but overlapping biological functions, and L-MYC overexpression has been validated to facilitate adult mouse MG reprogramming in retinal cultures[30]. Consistent with these findings, our study revealed that TRULI and CHIR combinatorial treatment markedly elevated both c-MYC and N-MYC protein levels in parallel with MG proliferation and reprogramming (**Figure** 7**E**-**G**). Furthermore, pharmacological suppression of MYC transcriptional activity via MYCi975, which disrupts MYC/MAX dimerization and abolishes MYC-dependent downstream signaling, nearly completely abrogated TRULI-and-CHIR-induced MG proliferation (**Figure** 7**H** and **I**). These results established MYC as an indispensable transcriptional regulator governing mammalian MG proliferative activation. Although previous studies have demonstrated overlapping functions of MYC in cell cycle control, pluripotency maintenance, and cellular reprogramming [27], our current work primarily focused on its pro-proliferative role in MG. Whether MYC additionally modulates MG reprogramming and neuronal fate transition requires further systematic investigation.

Precise cell cycle progression is orchestrated by coordinated activation of cyclin-dependent kinases (CDKs) and cyclin family proteins [31]. All retinal neurons in adult mammals are terminally differentiated and permanently exit the cell cycle; thus, cell cycle regulatory mechanisms in retinal tissues are predominantly characterized in proliferative retinal progenitor cells (RPCs). Cyclin D1, conventionally recognized as a core G1-phase regulator, exhibits broad expression across multiple cell cycle stages in RPCs[32, 33]. Genetic ablation of *Ccnd1* causes severe retinal hypoplasia and layered structural defects due to impaired RPC proliferation, highlighting its indispensable role in sustaining retinal progenitor expansion[34, 35]. Recent mechanistic studies further identified Cyclin D1 as a canonical downstream effector of Hippo-YAP signaling, wherein YAP activation bypasses Hippo-mediated suppression to trigger Cyclin D1-dependent MG proliferation[12, 13]. In line with these reports, our study confirmed that TRULI-induced Hippo inhibition upregulated Cyclin D1 expression, and blockade of YAP/β-catenin or MYC signaling reversed this inductive effect and abolished MG proliferation (**Figure** 6 and 7). These findings support a conserved regulatory axis whereby MYC bridges upstream YAP activation and downstream Cyclin D1-driven cell cycle progression in adult MG.

Cyclin D1-mediated cell cycle progression is tightly counterbalanced by CDK inhibitors, which ensure proper cell cycle exit, terminal differentiation, and retinal tissue morphogenesis[36]. Two canonical CDK inhibitors, p27^Kip1^ and p57^Kip2^, govern cell cycle withdrawal in distinct RPC subpopulations[36–38]. Specifically, p27^Kip1^ overexpression drives premature cell cycle exit in late-stage multipotent RPCs, facilitating their differentiation into bipolar cells, photoreceptors, and resident MG. Genetic rescue experiments further verified that loss of p27^Kip1^ rescues the proliferative defects of *Ccnd1*-deficient retinas, demonstrating cooperative and compensatory interactions between Cyclin D1 and p27^Kip1^ in retinal cell cycle regulation[39]. In the mature mammalian retina, p27^Kip1^ is constitutively expressed in quiescent MG, and its downregulation is a prerequisite for injury-triggered MG cell cycle re-entry [2]. Consistent with this model, targeted AAV-mediated Cyclin D1 overexpression combined with p27^Kip1^ knockdown is sufficient to induce robust proliferation in uninjured adult mouse MG [14]. Our results corroborated these observations, showing significant p27^Kip1^ protein downregulation together with Cyclin D1 upregulation in TRULI-and-CHIR-induced proliferative MG (**Figure** 6**E-H**). Furthermore, our study revealed a non-transcriptional regulatory mechanism underlying p27^Kip1^ suppression. Pharmacological inhibition of YAP, β-catenin, or MYC transcriptional activity further reduced p27^Kip1^ protein abundance (**Figure** 6**I** and **K**; **Figure** 7**J** and **M**), indicating that TRULI-and-CHIR-induced p27^Kip1^ downregulation occurred primarily at the post-translational level. Accumulating evidence suggests that p27^Kip1^ stability is controlled by Skp2-mediated ubiquitination and proteasomal degradation[40], and Akt-dependent Skp2 acetylation modulates p27^Kip1^ accumulation downstream of YAP signaling [41]. Our preliminary data show reduced Akt activity upon TRULI and CHIR combinatorial treatment, implying that enhanced Skp2-dependent proteasomal degradation may account for p27^Kip1^ downregulation in our system. However, retinal explant cultures exhibit high sensitivity to proteasome and Akt inhibitors (MG-132 and MK-2206, respectively), which prevented experimental validation of this post-translational regulatory mechanism. Future optimized culture systems or transgenic manipulations are required to dissect the precise post-transcriptional modulation of p27^Kip1^ in mammalian MG.

Hippo and Wnt signaling pathways are core modulators of cellular proliferation in regenerative biology. Consistently, previous genetic studies have independently verified that YAP [12, 13] and β-catenin[6] activation promote adult MG proliferation. Nevertheless, our study identified a robust synergistic pro-proliferative effect upon dual pathway co-activation, which cannot be recapitulated by single-pathway modulation alone. The molecular crosstalk between Hippo/YAP and Wnt/β-catenin signaling has been well documented in multiple biological contexts. In cardiomyocyte regeneration, nuclear YAP and β-catenin form functional complexes to cooperatively activate the reprogramming genes *Sox2* and *Snail2*[42]. Similarly, YAP/β-catenin transcriptional complexes drive proliferation, malignant transformation, and survival in multiple cancer types [43–45]. Several lines of evidence from our study supported the formation of YAP/β-catenin transcriptional complexes in activated MG. First, specific inhibition of either YAP-TEAD or β-catenin/TCF signaling abolished TRULI-and-CHIR-induced MG proliferation (**Figure** 5**F** and **G**). Second, although TRULI but not CHIR alone induced Cyclin D1 upregulation, both Verteporfin and ICG-001 effectively block Cyclin D1 induction under TRULI and CHIR treatment (**Figure** 6**I** and **J**). Third, TRULI and CHIR co-treatment significantly upregulates both canonical YAP target genes (*Bmp4*, *Cyr61*, *Ctgf*) and Wnt target genes (*Dkk1*, *Axin2*, *Lef1*), and inhibition of either pathway mediated transcription induced highly consistent target gene alteration patterns (**Figure S**6**A**-**F**). Collectively, these data suggest that YAP and β-catenin may assemble transcriptional complexes to drive robust MG proliferation, though the detailed structural and regulatory mechanisms require further exploration.

In zebrafish, MG proliferation and progenitor reprogramming are tightly coupled processes during retinal regeneration[1]. Similarly, YAP5SA-mediated Hippo bypass in murine MG drives cell cycle re-entry accompanied by RPC-like reprogramming[13]. Our study confirmed that TRULI-and-CHIR-induced proliferative MG upregulated the core progenitor marker PAX6 and multiple RPC signature genes (**Figure** 3**A**-**E**, **Figure S**3**B**), indicating successful glia-to-progenitor conversion. Intriguingly, PAX6 upregulation was not further enhanced under enhanced MG proliferation triggered by TRULI and CHIR treatment, with comparable PAX6 levels between the control and combinatorial groups (**Figure** 3**D**). This observation suggests that MG cell cycle re-entry and RPC gene reactivation regulatory machineries may be partially uncoupled in the mammalian system, implying that progenitor-like reprogramming may be attributed to intrinsic retinal explant injury rather than proliferation.

The neurogenic potential of reprogrammed MG is a critical determinant of functional retinal regeneration and remains a major bottleneck in mammalian regenerative research. Constitutive YAP activation often locks MG in a proliferative state with minimal neuronal differentiation capacity. Recent studies using transient Hippo-resistant YAP variants in adult mouse MG suggested that most of the proliferative MG persistently retain glial identity, with sustained SOX9 expression and negligible induction of bipolar cell markers[46]. In agreement with these findings, our study showed that nearly all proliferative MG after TRULI and CHIR combinatorial treatment maintained SOX9 expression (**Figure S**3**F**), further supporting the notion that the regulatory mechanisms governing proliferation, RPC formation, and neuronal trans-differentiation are at least partially uncoupled in mammalian MG. Thus, unlocking the neurogenic potential of MG remains the most pressing open question in MG-mediated retinal regeneration. One established inhibitory mechanism is persistent Notch signaling activity in adult MG, which blocks glia-to-neuron conversion[7, 8, 47]. In addition, *Ascl1*, a critical proneural factor required for zebrafish MG reprogramming and mammalian retinal neuron development[48, 49], remains largely silenced in MG (**Figure S**3**B**), which may account for the limited neuronal trans-differentiation potential observed in mammalian MG following combined TRULI and CHIR treatment.

Despite this limited neuronal differentiation potential, we observed specific upregulation of Otx2, an early transcription factor governing bipolar and photoreceptor lineage specification, in proliferative MG (**Figure** 4**A** and **B**). In contrast, no amacrine or ganglion cell markers were detected within our experimental time window (**Figure S**4). These results demonstrate that chemically reprogrammed MG possess partial yet restricted neurogenic potential, with a preferential bias toward the bipolar/photoreceptor lineage. In our explant model, PAX6 (**Figure** 3) and its downstream proneural bHLH transcription factor Neurogenin2 (Neurog2) (**Figure S**3**B**) were upregulated. During mouse retinal development, Neurog2 cooperates with Ascl1 to regulate Otx2 expression and thereby promotes bipolar/photoreceptor genesis [49, 50]. Moreover, CHX10 is a key transcription factor governing fate specification between bipolar cells and rod photoreceptors[51]. Since CHX10 was not induced in MG after combined TRULI and CHIR treatment (**Figure** 3**G**), reprogramming MG under these conditions may exhibit a stronger preference toward the rod photoreceptor fate. Indeed, we observed tdTomato+ MG that very occasionally expressed mature photoreceptor markers recoverin or SAG (**Figure** 4**C** and **D**). Whether this lineage preference arises from developmental homology between MG and inner retinal neurons or microenvironmental constraints of the retinal explant model remains to be elucidated.

The present study has several obvious limitations. All functional and molecular characterizations were performed in *ex vivo* retinal explant cultures, which lack intact vascular circulation, immune microenvironment, and physiological intraocular drug metabolism. These deficiencies may alter MG behaviors and pathway activation dynamics compared with the *in vivo* scenario. In addition, our preliminary data indicated that both TRULI and CHIR exhibited poor blood–retina barrier permeability, limiting the efficacy of systemic intraperitoneal drug administration. Future studies will focus on optimizing local intraocular delivery strategies and validating the regenerative efficacy and safety of this chemical combination *in vivo* to advance its translational potential for retinal degenerative disease treatment.

### Data availability

Raw RNA-seq data have been deposited in the NCBI Sequence Read Archive (SRA) under BioProject accession PRJNA1521908. Processed expression matrices will be deposited in the NCBI Gene Expression Omnibus (GEO) before acceptance; GEO accession number GSE348108. Uncropped Western blot exposures will be made available with the submission. The Majorbio cloud project used for internal analysis is access restricted and does not replace deposition in a public repository. Other data are available from the corresponding author upon reasonable request.

## Acknowledgements

This work was supported by grants from the Natural Science Foundation of Fujian Province (Grant No. 2026J001012, 2026J0011725), Natural Science Foundation of Xiamen, China (Grant No.3502Z202373024). The authors thank Prof. Wenjun Xiong from City University of Hong Kong and Prof. Yuanyuan Chen from University of Pittsburg for insightful suggestions and technical support, Jingru Huang and Xiang You from Central Lab, School of Medicine, Xiamen University for technical support in confocal imaging.

## Author contributions

**Wanling He**: Conceptualization, Methodology, Investigation, Data curation, Formal analysis, Visualization, Writing – original draft. **Yaqiong Yang**: Conceptualization, Methodology, Investigation. **Ziyi Sha**: Investigation. **Qinrui Hu**: Funding acquisition. **Xiying Zhai**: Investigation. **Jintao Shi**: Resources. **Lifan Yang**: Resources. **Zihan Guo**: Resources. **Mouxin Zhang**: Resources. **Yuehua Zhang**: Resources. **Wei Li**: Methodology. **Liying Tang**: Funding acquisition, Data curation. **Yi Liao**: Conceptualization, Supervision, Project administration, Funding acquisition, Writing – review & editing. All authors read and approved the final manuscript.

