## supplemental figures for "Pharmacological co-targeting of Hippo and Wnt pathways induces robust Müller glia proliferation and reprogramming in the adult mouse retina"

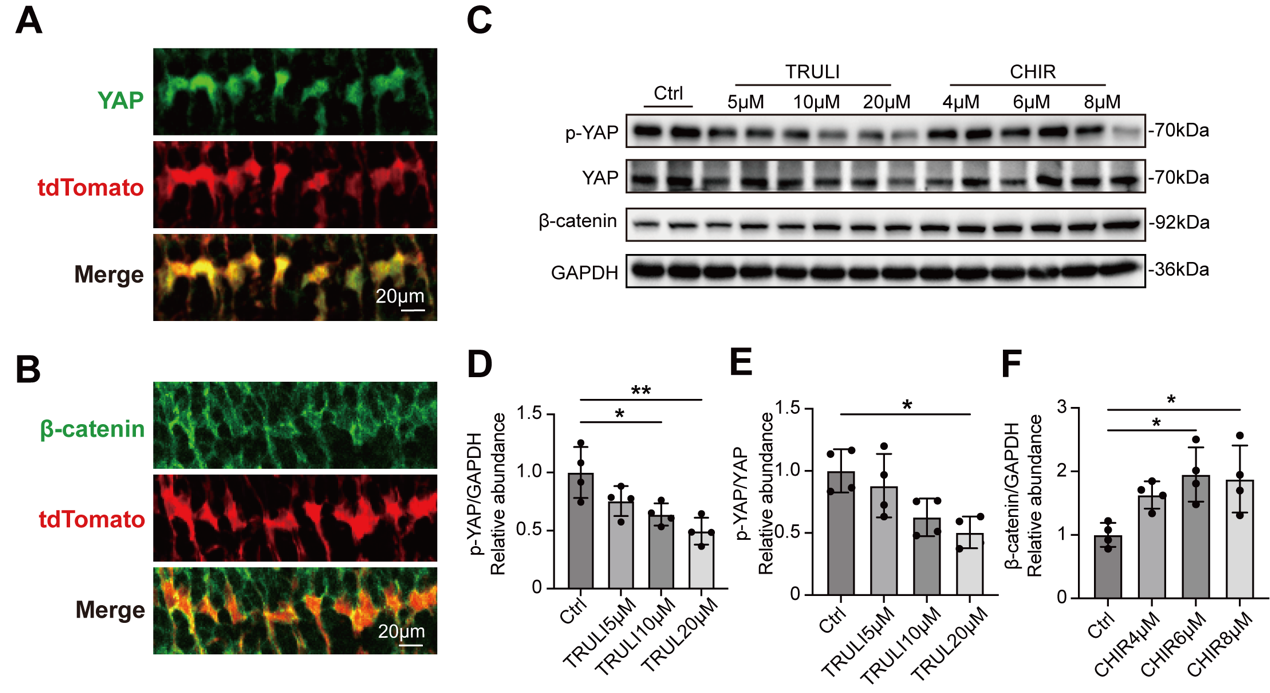

**Figure S1.** The regulatory effects of TRULI and CHIR on the Hippo and Wnt signaling pathways. (**A**) Immunofluorescence staining for YAP (*green*) in Glast-Cre^ERT2^; tdTomato (*red*) labelled Müller glia (MG) in fresh retinal sections. (**B**) Immunofluorescence of β-catenin (*green*) in Glast-Cre^ERT2^; tdTomato (*red*) labelled MG in retinal sections of fresh retina. (**C**) Western blot analysis of p-YAP, total YAP, and β-catenin protein levels in retinal explants after 3 days of treatment with different concentrations of TRULI or CHIR. GAPDH was used as a loading control. (D–F) Quantification of p-YAP protein levels normalized to GAPDH (D), the p-YAP/YAP ratio (E), and β-catenin protein levels normalized to GAPDH (F), corresponding to the blots in panel (C). n=4 retinas per group. Data are presented as mean ± SD; one-way ANOVA with Tukey's multiple-comparisons test. **p*<0.05, ***p*<0.01. Scale bar: 20 μm.

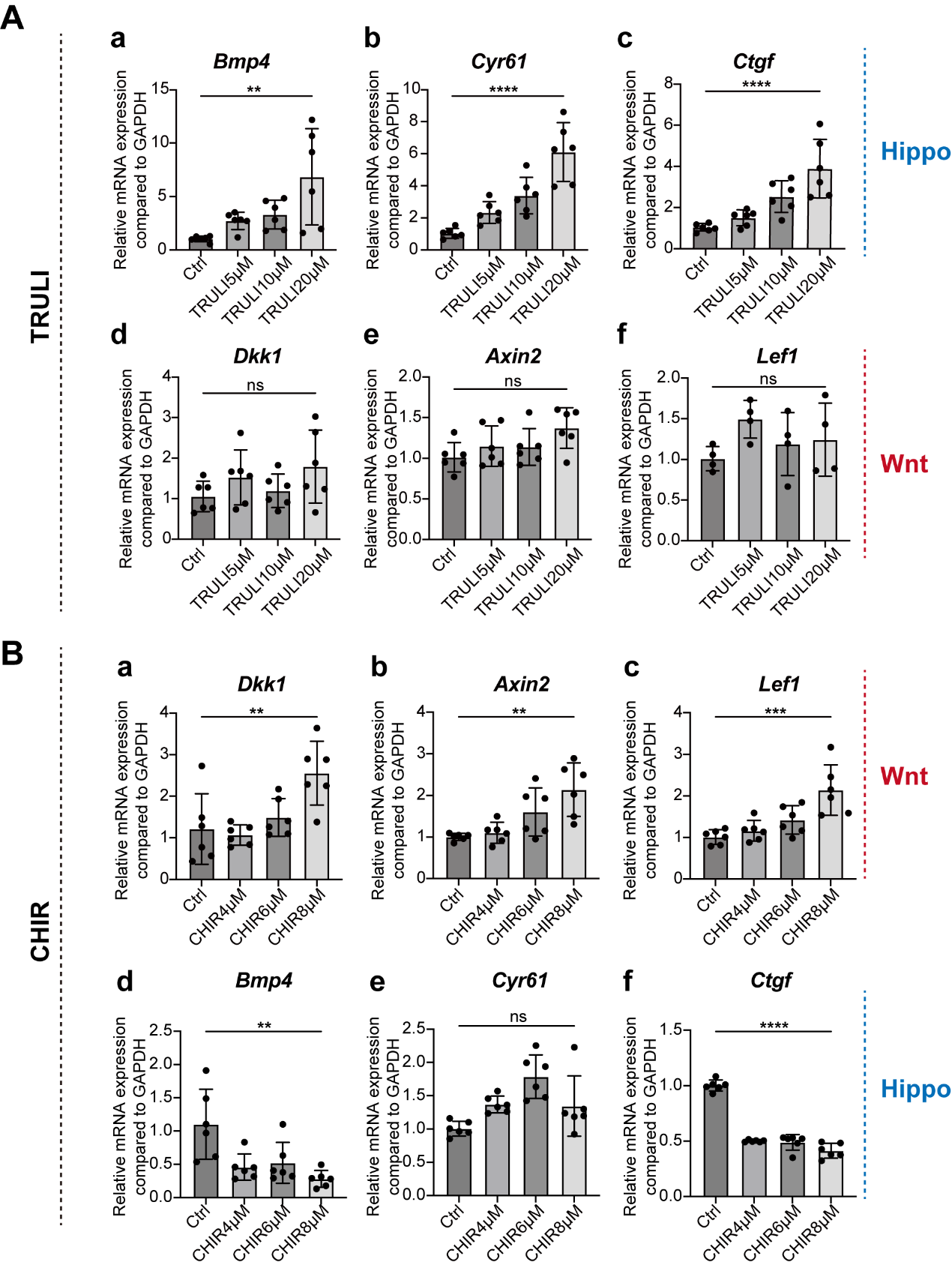

**Figure S2.** TRULI and CHIR dose-dependently regulate Hippo and Wnt pathway target genes. (**A**) qRT-PCR analysis of mRNA levels of YAP target genes *Bmp4*, *Cyr61,* and *Ctgf* and Wnt target genes *Dkk1*, *Axin2*, and *Lef1* after treatment with different concentrations of TRULI. (**B**) qRT-PCR analysis of mRNA levels of Wnt target genes *Dkk1*, *Axin2*, and *Lef1* and Hippo target genes *Bmp4*, *Cyr61,* and *Ctgf* after treatment with different concentrations of CHIR. n=6 retinas per group. Data are presented as mean ± SD; one-way ANOVA with Tukey's multiple-comparisons test. ***p*<0.01, ****p*<0.001, **** *p*<0.0001, ns, not significant.

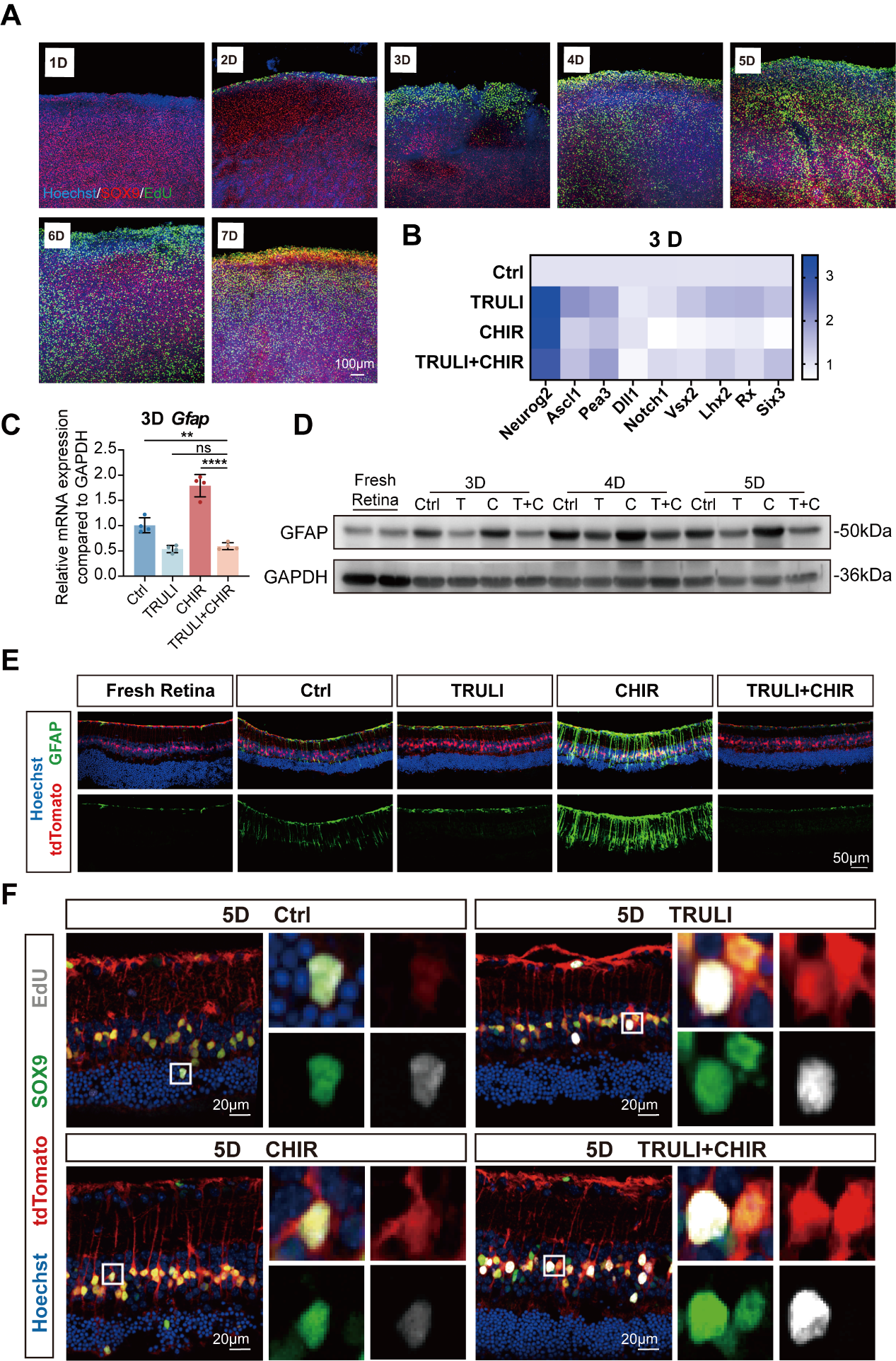

**Figure S3.** The trans-differentiation fate of MG after TRULI and CHIR treatment. (A) Immunofluorescence staining for SOX9 (*red*) and EdU incorporation (*green*) in the peripheral regions of retinal explants after TRULI+CHIR treatment with continuous EdU labelling for 1-7 consecutive days. Cell nuclei were counterstained with Hoechst (*blue*). (**B**) Heatmap showing the mRNA expression profiles of progenitor-associated genes in retinal tissues after 3 days of treatment with vehicle control (Ctrl), TRULI, CHIR, or TRULI+CHIR treatment; n=4 retinas per group. (**C**) qRT-PCR analysis of *Gfap* mRNA levels after 3 days of treatment with Ctrl, TRULI, CHIR, or TRULI+CHIR; n=4 retinas per group. (**D**) Western blot analysis of GFAP protein expression in fresh retinas and retinal explants after 3-5 days of Ctrl, TRULI (T), CHIR (C), or TRULI+CHIR (T+C) treatment. GAPDH was used as an internal loading control. (**E**) Immunofluorescence staining for GFAP (*green*) in Glast-Cre^ERT2^; tdTomato (*red*)- labelled MG in sections of fresh retina and retinal explants after 5 days of Ctrl, TRULI, CHIR, or TRULI+CHIR treatment. Cell nuclei were counterstained with Hoechst (*blue*). (**F**) Immunofluorescence staining for SOX9 (*green*) and EdU incorporation (*gray*) in Glast-Cre^ERT2^; tdTomato (*red*) -labelled MG in retinal sections after 5 days of Ctrl, TRULI, CHIR, or TRULI+CHIR treatment. Cell nuclei were counterstained with Hoechst (*blue*). Data are presented as mean ± SD; one-way ANOVA with Tukey's multiple-comparisons test. ***p*<0.01, **** *p*<0.0001, ns, not significant. Scale bars: 100 μm in (**A**); 50 μm in (**E**), and 20μm in (**F**), as indicated.

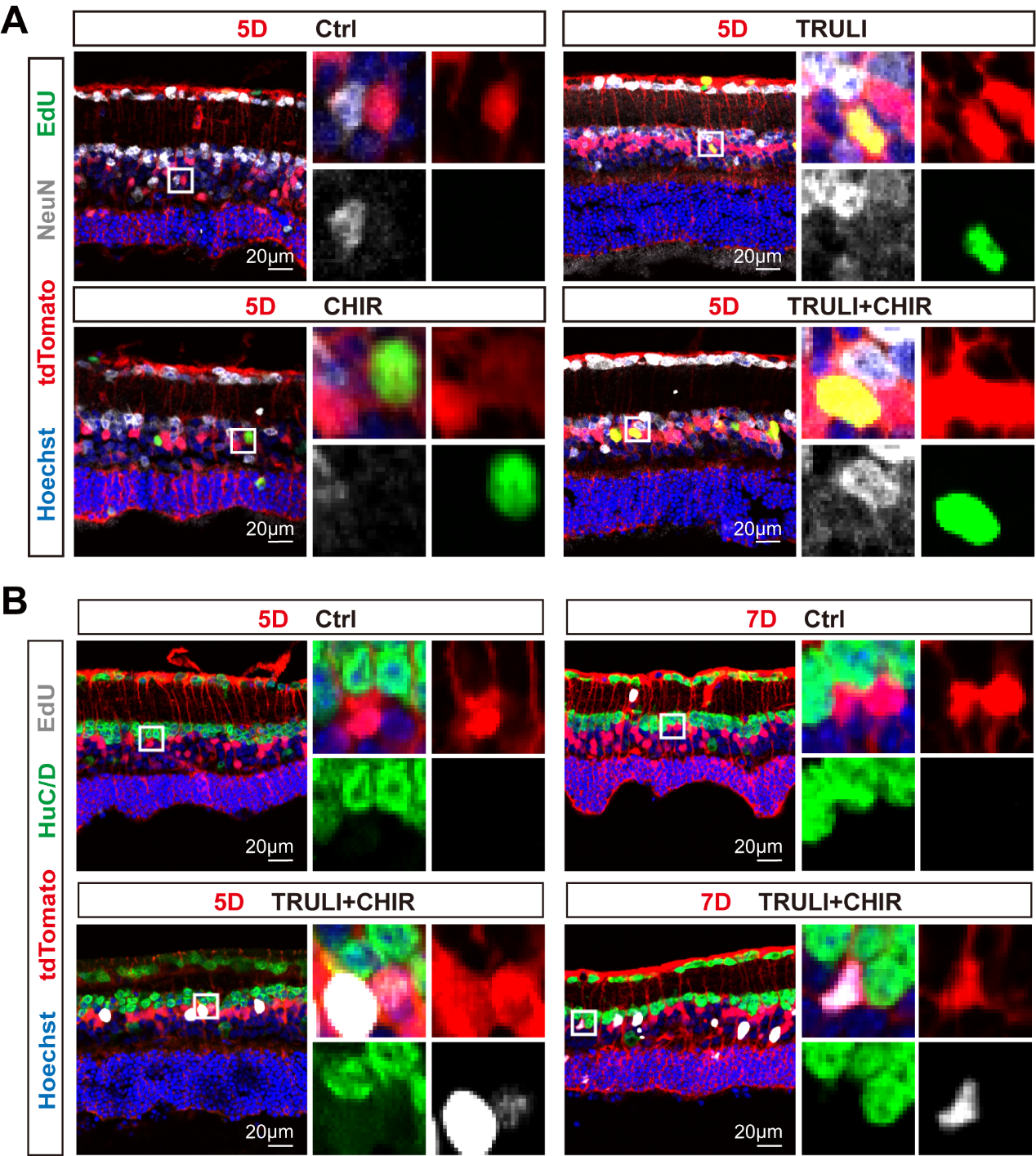

**Figure S4.** TRULI and CHIR treatment does not induce the expression of amacrine or ganglion cell lineage markers in MG. (**A**) Immunofluorescence staining for NeuN (*gray*) and EdU incorporation (*green*) in Glast-Cre^ERT2^; tdTomato (*red*)-labelled MG in retinal sections after 5 days of Ctrl, TRULI, CHIR, or TRULI+CHIR treatment. Cell nuclei were counterstained with Hoechst (*blue*). (B) Immunofluorescence staining for HuC/D (*green*) and EdU incorporation (*gray*) in Glast-Cre^ERT2^; tdTomato (*red*) -labelled MG in retinal sections after 5 and 7 days of Ctrl or TRULI+CHIR treatment. Cell nuclei were counterstained with Hoechst (*blue*). Scale bars: 20 μm.

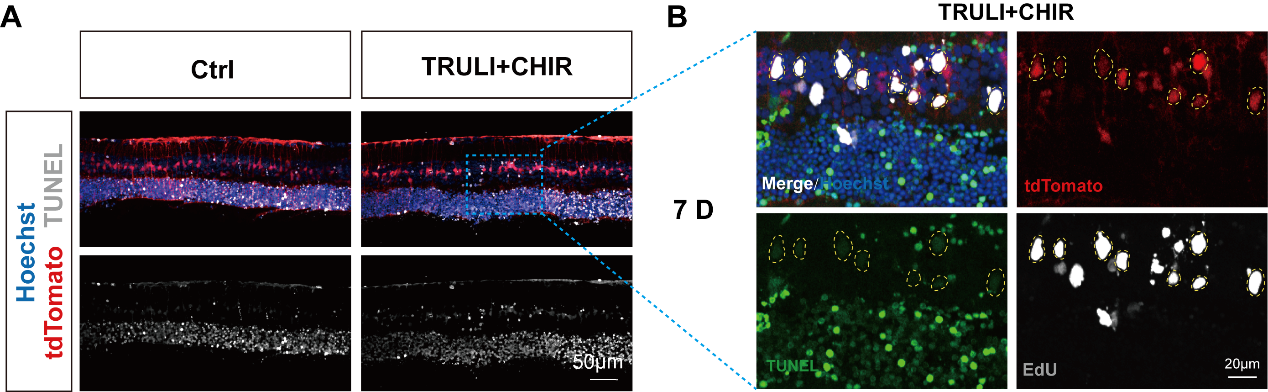

**Figure S5.** TUNEL staining analysis of retinal explants on day 7. (**A**) Immunofluorescence staining for TUNEL signals (*grey*) in Glast-Cre^ERT2^; tdTomato (*red*) -labelled MG in retinal sections after 7 days of Ctrl and TRULI+CHIR treatment. Cell nuclei were counterstained with Hoechst (*blue*). (**B**) Magnified view showing co-staining of TUNEL (*green*) and EdU (*grey*) in Glast-Cre^ERT2^; tdTomato (*red*) -labelled MG from the TRULI+CHIR group at day 7. Scale bars: 50 μm in (**A**); 20 μm in (**B**).

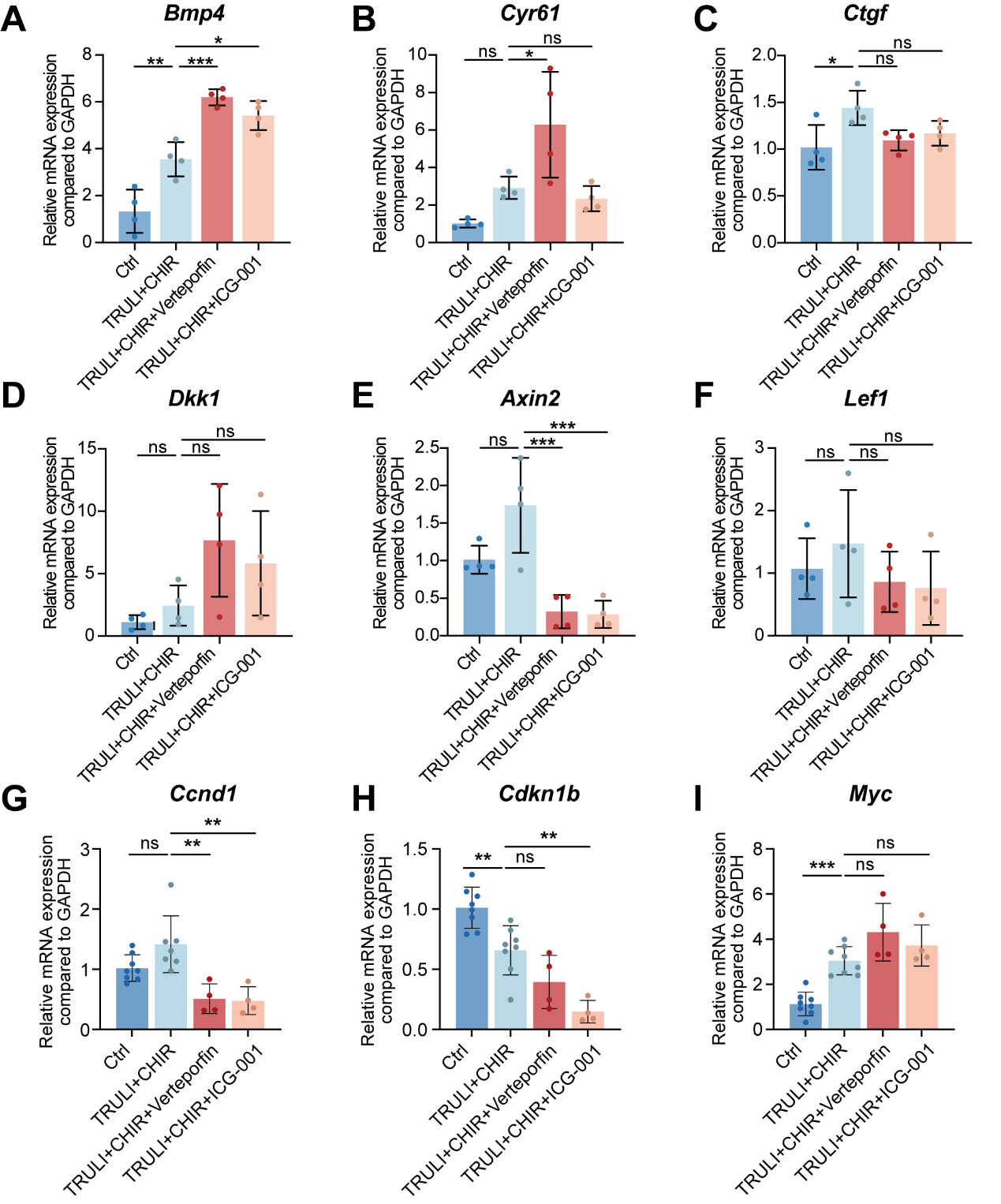

**Figure S6.** Effects of Verteporfin or ICG-001 on the expression of target genes after TRULI and CHIR treatment. (**A**–**C**) qRT-PCR analysis of mRNA levels of Hippo target genes *Bmp4*, *Cyr61*, and *Ctgf* after 4 days of Ctrl, TRULI+CHIR, TRULI+CHIR with Verteporfin or TRULI+CHIR with ICG-001 treatment. (**D**-**F**) qRT-PCR quantification of mRNA levels of Wnt target genes *Dkk1*, *Axin2*, and *Lef1* after 4 days of Ctrl, TRULI+CHIR, TRULI+CHIR with Verteporfin or TRULI+CHIR with ICG-001 treatment. (**G**-**I**) qRT-PCR quantification of mRNA levels of cell cycle regulatory genes *Ccnd1*, *Cdkn1b*, and *Myc* after 4 days of Ctrl, TRULI+CHIR, TRULI+CHIR with Verteporfin or TRULI+CHIR with ICG-001 treatment. n≥4 retinas per group. Data are presented as mean ± SD; one-way ANOVA with Tukey's multiple-comparisons test. **p*<0.05, ***p*<0.01, *** *p*<0.001, ns, not significant.

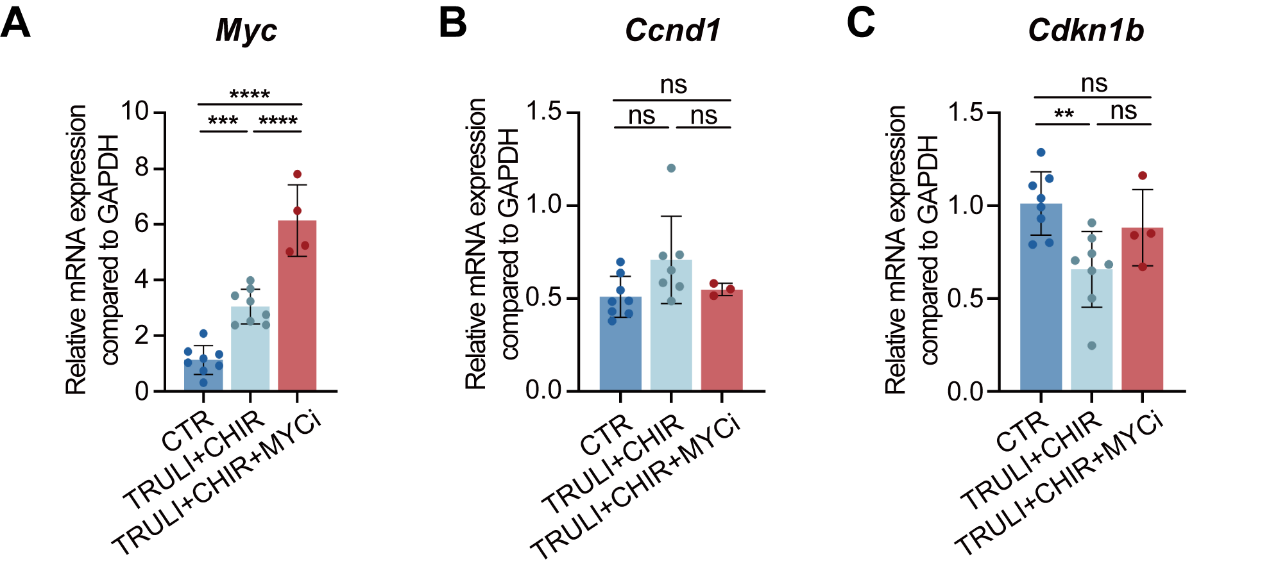

**Figure S7.** Effects of MYC inhibitor MYCi975 on the expression of cell cycle regulatory genes. qRT-PCR quantification of mRNA levels of *Myc* (**A**), *Ccnd1*(**B**), and *Cdkn1b*(**C**) after 4 days of Ctrl, TRULI+CHIR, or TRULI+CHIR with MYCi975 treatment. n≥3 retinas per group. Data are presented as mean ± SD; one-way ANOVA with Tukey's multiple-comparisons test. **p*<0.05, ***p*<0.01, ****p*<0.001, *****p*<0.0001; ns, not significant.

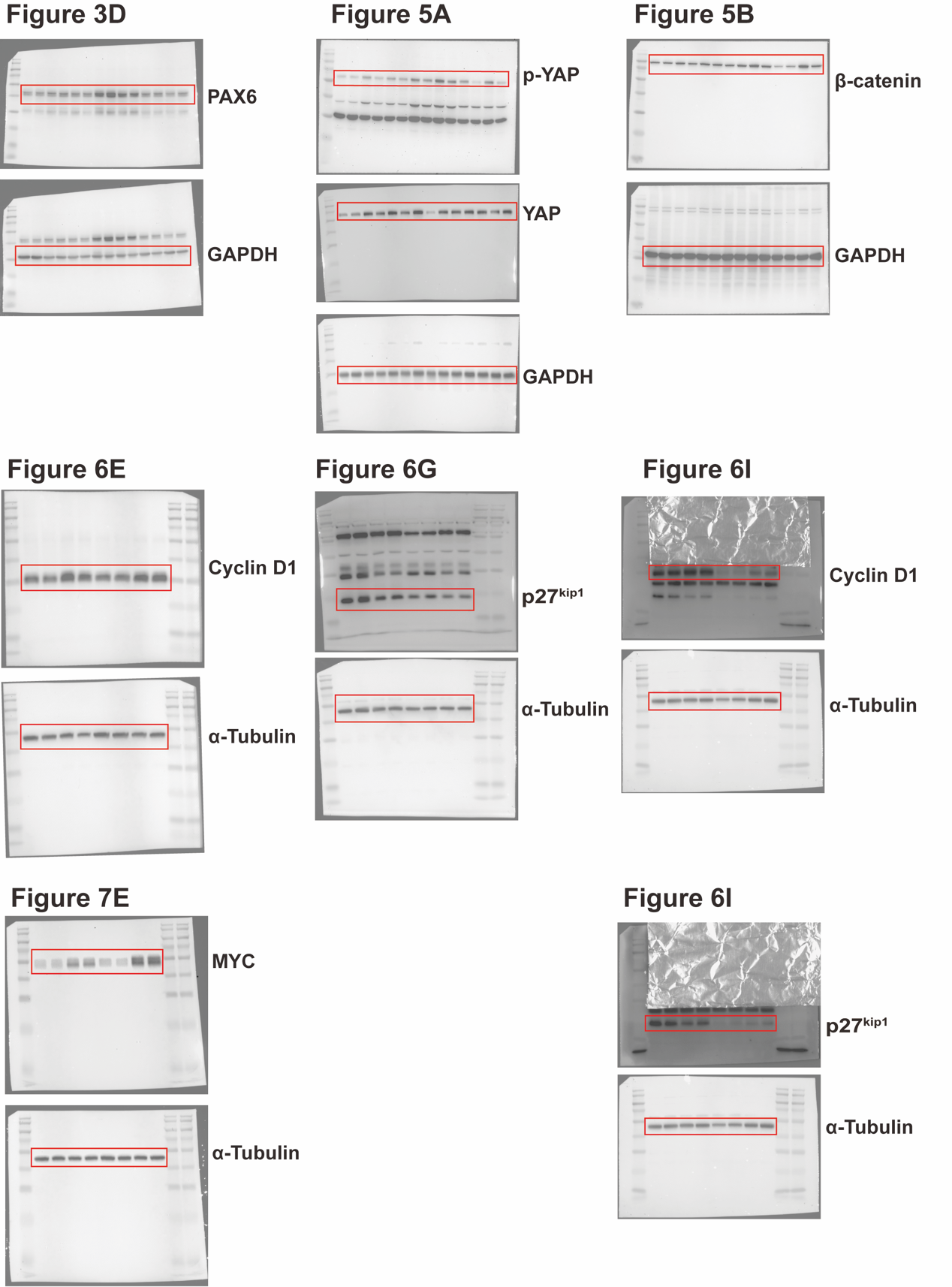

**Figure S8**. Original blots for Western blot images presented in manuscript (group 1).

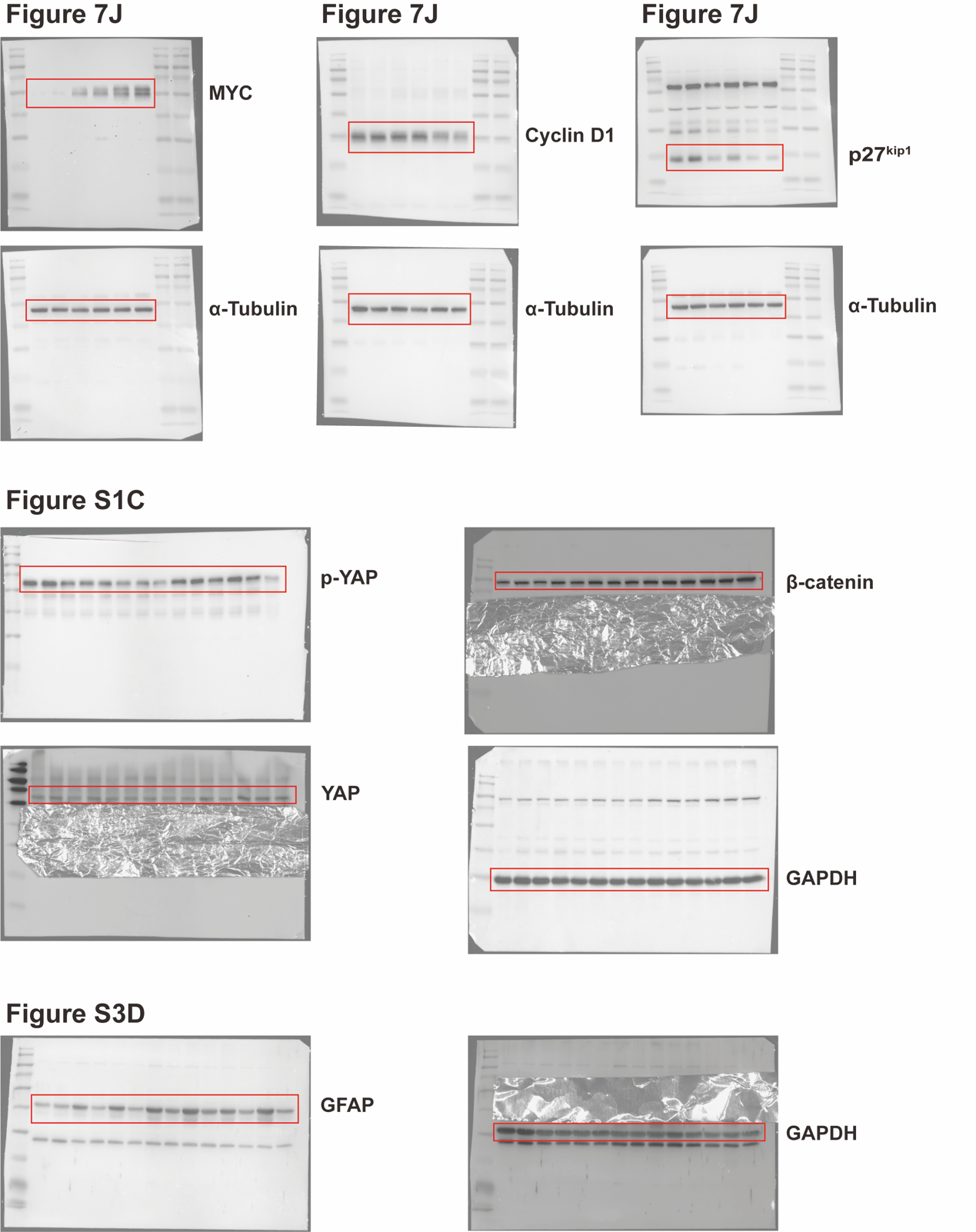

**Figure S9**. Original blots for Western blot images presented in manuscript (group 2).

**Supplementary Tables**

**Supplementary Table** 1| Antibodies used in this study

| **Target** | **Antibody name** | **Manufacturer** | **RRID** | **Catalog no.** | **Host**  **/type** | **Application**  **/dilution** |
| --- | --- | --- | --- | --- | --- | --- |
| β-Catenin | β-Catenin (D10A8) XP® Rabbit mAb | Cell Signaling Technology | AB_11127855 | 8480T | Rabbit monoclonal | IF 1:100;  WB 1:1000 |
| CHX10/VSX2 | CHX10/VSX2 Antibody | Novus Biologicals | AB_11022841 | NBP1-84476 | Rabbit polyclonal | IF 1:200 |
| HuC/D | HuD + HuC (EPR19098) | Abcam | AB_2864321 | ab184267 | Rabbit monoclonal | IF 1:500 |
| SOX9 | SOX9 (EPR14335) | Abcam | AB_2715497 | ab185230 | Rabbit monoclonal | IF 1:500 |
| GFAP | GFAP Rabbit mAb (ARC76397) | ABclonal | AB_2862549 | A27997 | Rabbit monoclonal | IF 1:300;  WB 1:5000 |
| Ki67 | Ki67 (B56) | Abcam | AB_2934265 | ab279653 | Mouse monoclonal | IF 1:500 |
| Cyclin D1 | [KO Validated] Cyclin D1 Rabbit mAb | ABclonal | AB_2862530 | A19038 | Rabbit monoclonal | WB 1:1000 |
| PAX6 | PAX6 Rabbit pAb | ABclonal | AB_2767871 | A7334 | Rabbit polyclonal | WB 1:1000 |
| PAX6 | Pax-6 (PAX6) | Santa Cruz Biotechnology | AB_1127044 | sc-81649 | Mouse monoclonal IgG1 | IF 1:200 |
| p27Kip1 | CDKN1B/p27KIP1 Rabbit pAb | ABclonal | AB_2757102 | A0290 | Rabbit polyclonal | WB 1:2000 |
| NeuN | NeuN (EPR12763) | Abcam | AB_2532109 | ab177487 | Rabbit monoclonal | IF 1:500 |
| c-Myc/N-Myc | c-Myc/N-Myc (D3N8F) Rabbit mAb | Cell Signaling Technology | AB_2631168 | 13987S | Rabbit monoclonal | IF 1:200;  WB 1:1000 |
| Recoverin | RCVRN/Recoverin Rabbit pAb | ABclonal | AB_2767006 | A6404 | Rabbit polyclonal | IF 1:300 |
| SAG | SAG Rabbit pAb | ABclonal | AB_2759893 | A13045 | Rabbit polyclonal | IF 1:200 |
| α-Tubulin | α-Tubulin (1E4C11) Mouse mAb | Proteintech | AB_11042766 | 66031-1-Ig | Mouse monoclonal | WB 1:5000 |
| GAPDH | GAPDH Rabbit mAb (ARC50882) | ABclonal | AB_3754674 | A22420 | Rabbit monoclonal | WB 1:5000 |
| p-YAP (Ser127) | Phospho-YAP (Ser127) Antibody | Cell Signaling Technology | AB_2218913 | 4911S | Rabbit polyclonal | WB 1:1000 |
| YAP1 | [KO Validated] YAP1 Rabbit mAb | ABclonal | AB_2862627 | A19134 | Rabbit monoclonal | IF 1:500;  WB 1:1000 |
| OTX2 | Human Otx2 Antibody | R&D Systems (Bio-Techne) | AB_2157172 | AF1979 | Goat polyclonal | IF 10 μg/mL |
| Fluorescent secondary antibody | ABflo® 488-conjugated Goat anti-Rabbit IgG (H+L) | ABclonal | AB_2768317 | AS073 | Goat; affinity-purified | IF 1:300 |
| Fluorescent secondary antibody | ABflo® 488-conjugated Goat anti-Mouse IgG (H+L) | ABclonal | AB_2768316 | AS076 | Goat; affinity-purified | IF 1:300 |
| Fluorescent secondary antibody | ABflo® 647-conjugated Goat anti-Rabbit IgG (H+L) | ABclonal | AB_2768330 | AS060 | Goat; affinity-purified | IF 1:300 |
| Fluorescent secondary antibody | ABflo® 647-conjugated Goat anti-Mouse IgG (H+L) | ABclonal | AB_2768329 | AS059 | Goat; affinity-purified | IF 1:300 |
| Fluorescent secondary antibody | FITC-conjugated Donkey anti-Goat IgG (H+L) | ABclonal | AB_2769473 | AS032 | Donkey; affinity-purified | IF 1:300 |
| HRP-conjugated secondary antibody | Goat Anti-Rabbit IgG (H+L)-HRP Conjugate | Bio-Rad | AB_11125142 | 1706515 | Goat polyclonal secondary | WB 1:5000 |
| HRP-conjugated secondary antibody | Goat Anti-Mouse IgG (H+L)-HRP Conjugate | Bio-Rad | AB_11125547 | 1706516 | Goat polyclonal secondary | WB 1:5000 |

**Supplementary Table** 2| qRT-PCR primer sequences

| **Gene** | **Forward primer (5′–3′)** | **Reverse primer (5′–3′)** |
| --- | --- | --- |
| *Dkk1* | TACCCTTGCGCTGAAGATGA | TGGTCAGAGGGCATGCATAT |
| *Bmp4* | TCCTGGTAACCGAATGCTGA | CTGTAGAAGTGTCGCCTCGA |
| *Cyr61* | AACGAGGACTGCAGCAAAAC | GCGCCATCAATACATGTGCA |
| *Ctgf* | GAGCGCCTGTTCTAAGACCT | GGCTTGGCGATTTTAGGTGT |
| *Axin2* | AGAAGAGGAGTGGACGTGTG | AGACATAGCCGGAACCTACG |
| *Ccnd1* | GCGGATGAGAACAAGCAGAC | AGGAGAGGAAGTTGTTGGGG |
| *Gapdh* | TGGAAAGCTGTGGCGTGAT | GGGTAGGAACACGGAAGGC |
| *Lef1* | AGGAGCCCAAAAGACCTCAT | GCTGTCTCTCTTTCCGTGCT |
| *Gfap* | GCCACCAGTAACATGCAAGA | GCTCTAGGGACTCGTTCGTG |
| *Pax6* | CTGAGGAACCAGAGAAGACAGG | CATGGAACCTGATGTGAAGGAGG |
| *Myc* | TCTCCACTCACCAGCACAACTACG | ATCTGCTTCAGGACCCT |
| *Cdkn1b* | TTGGGTCTCAGGCAAACTCT | TCTTCTGTTCTGTTGGCCCT |
| *Mki67* | GATGGAAGCATTGTGAGAACCA | CCTGCTCTTCCACAGATTCAAG |

**Supplementary Table** 3| Genotyping primer sequences

| **Line** | **Primer** | **Sequence (5′–3′)** | **Purpose** | **Product** |
| --- | --- | --- | --- | --- |
| Glast-CreERT2 | 10110 | ACAATCTGGCCTGCTACCAAAGC | Transgene | ~600 bp |
| Glast-CreERT2 | 10112 | CCAGTGAAACAGCATTGCTGTC | Transgene | ~600 bp |
| Glast-CreERT2 | oIMR8744 | CAAATGTTGCTTGTCTGGTG | Internal control | 200 bp |
| Glast-CreERT2 | oIMR8745 | GTCAGTCGAGTGCACAGTTT | Internal control | 200 bp |
| Ai14 | oIMR9020 | AAGGGAGCTGCAGTGGAGTA | Wild-type F | 297 bp |
| Ai14 | oIMR9021 | CCGAAAATCTGTGGGAAGTC | Wild-type R | 297 bp |
| Ai14 | oIMR9103 | GGCATTAAAGCAGCGTATCC | Mutant R | ~200 bp |
| Ai14 | oIMR9105 | CTGTTCCTGTACGGCATGG | Mutant F | ~200 bp |
